# Programmable RNA Sensing for Cell Monitoring and Manipulation

**DOI:** 10.1101/2022.05.25.493141

**Authors:** Yongjun Qian, Jiayun Li, Shengli Zhao, Elizabeth Matthews, Michael Adoff, Weixin Zhong, Xu An, Michele Yeo, Christine Park, Bor-Shuen Wang, Derek Southwell, Z. Josh Huang

## Abstract

RNAs are the central and universal mediator of genetic information underlying the diversity of cell types and cell states, which together shape tissue organization and organismal function across species and life spans. Despite advances in RNA sequencing and massive accumulation of transcriptome datasets across life sciences, the dearth of technologies that leverage RNAs to observe and manipulate cell types remains a prohibitive bottleneck in biology and medicine. Here, we describe CellREADR (<u>Cell</u> access through <u>R</u>NA sensing by <u>E</u>ndogenous <u>A</u>DA<u>R)</u>, a programmable RNA sensing technology that leverages RNA editing mediated by ADAR (adenosine deaminase acting on RNA) for coupling the detection of cell-defining RNAs with translation of effector proteins. Viral delivery of CellREADR conferred specific cell type access in mouse and rat brains and in ex vivo human brain tissues. Furthermore, CellREADR enabled recording and control of neuron types in behaving mice. CellREADR thus highlights the potential for RNA-based monitoring and editing of animal cells in ways that are specific, versatile, easy, and generalizable across organ systems and species, with broad applications in biology, biotechnology, and programmable RNA medicine.

## INTRODUCTION

The diversity of cells underlies the diversity of life forms and of physiological systems within individual organisms^1^. Reading from a singular genome differentially packaged into numerous epigenomes, individual cells of the same organism generate distinct transcriptomes as RNA intermediates to implement cell specific phenotypes and physiology; thus RNAs are the central and universal messengers that convey customized genetic instructions in diverse and individual cell types and cell states that together shape tissue organization and system function across species and life spans. Indeed, all bodily functions emerge from interactions among cell types, and aberrant cell physiology and function give rise to myriad diseases^2^. Recent advances in single-cell RNA sequencing approaches promise to identify all molecularly-defined cell types in the human body and in many other organisms^3,4^. Beyond transcriptome profiling, it is necessary to monitor and manipulate each and every cell type in order to identify their specific roles in tissue architecture and system function within organisms; cell-specific monitoring and intervention is also key to precision diagnosis and treatment of many human diseases. To achieve these, we need cell type technologies that are specific, facile, scalable, economical, and general across species, akin to gene editing in genetics.

To date, almost all genetic approaches for accessing cell types rely on DNA-based transcriptional regulatory elements for expressing tool-genes (e.g. markers, sensors, effectors) that mimic certain cell-specific RNA expression, mostly through germline engineering in only a handful of organisms ^5–7^. However, all germline approaches, including those based on CRISPR ^8–10^, are inherently cumbersome, slow, difficult to scale and generalize, and raise ethical issues especially in primates and humans^11,12^. Recently, transcriptional enhancer-based viral vectors show promise for targeting cell types in animals^13–19^; but such enhancers are difficult to identify and validate, often requiring large-scale effort^13^ due to their complex relationship to target genes and cell types. Ultimately, all DNA- and transcription-based approaches are inherently indirect in attempting to mimic and leverage cell-specific RNA expression patterns.

Here, we report a new class of RNA sensing-dependent protein translation technology, bypassing the entire DNA-based transcriptional process, and apply it to genetic access of cell types. We invented CellREADR, <u>Cell</u> access through <u>R</u>NA sensing by <u>E</u>ndogenous <u>A</u>DA<u>R</u> (adenosine deaminase acting on RNA), which harnesses an RNA sensing and editing mechanism ubiquitous to all animal cells for detecting specific cellular RNAs then switching on translation of effector proteins to monitor and manipulate the cell. Deployable as a single RNA molecule operating through Watson-Crick base pairing, CellREADR as a cell engineering technology is inherently specific, easy, scalable, programmable, and general across species.

### CellREADR design and implementation in mammalian cells

RNA editing is a widespread and robust post-transcriptional mechanism essential to the metazoan gene regulatory toolkit implicated in recoding, splicing, microRNA targeting, and other RNA modulatory processes^20,21^. The most prevalent form of RNA editing is adenosine-to-inosine (A→I) conversion, catalyzed by ADARs; inosine is subsequently recognized as guanosine (G) by the cellular machinery^21^. ADARs recognize and are recruited by stretches of base-paired double-stranded (ds) RNAs^20^, and can therefore operate as a sequence-guided base editing machine, which can be harnessed for transcriptome editing^22–26^. As ADARs are ubiquitous in animal cells^27^, we designed CellREADR as a single and modular readrRNA molecule (**Fig. 1a**). The 5’ region of readrRNA contains a sensor domain of ∼300 nucleotides, which is complementary to and thus detects a specific cellular RNA through sequence-specific base pairing. This sensor domain contains one or more ADAR-editable STOP codons that act as a translation switch; we name this region the sense-edit-switch RNA (sesRNA). Downstream to the sesRNA and in-frame to the STOP is sequence coding for the self-cleaving peptide T2A, followed in-frame by an effector RNA (efRNA) region coding for various effector proteins. The entire readrRNA is several kilobases, depending on which specific sensors and effectors are included, and thus is deliverable to cells through viral vectors or liposome nanoparticles. In cells expressing the target RNA, sesRNA forms a dsRNA with the target, which recruits ADARs to assemble an editing complex. At the editable STOP codon, ADARs convert A to I, which pairs with the opposing C in the target RNA. This A→G substitution converts a UAG STOP codon to a UI(G)G tryptophan codon, switching on translation of the efRNA. The in-frame translation generates a fusion protein comprising an N-terminal peptide, T2A, and C-terminal effector, which then self-cleaves through T2A and releases the functional effector protein (**Fig. 1a**). readrRNA is expected to remain inert in cells that do not express the target RNA.

**Fig. 1:**
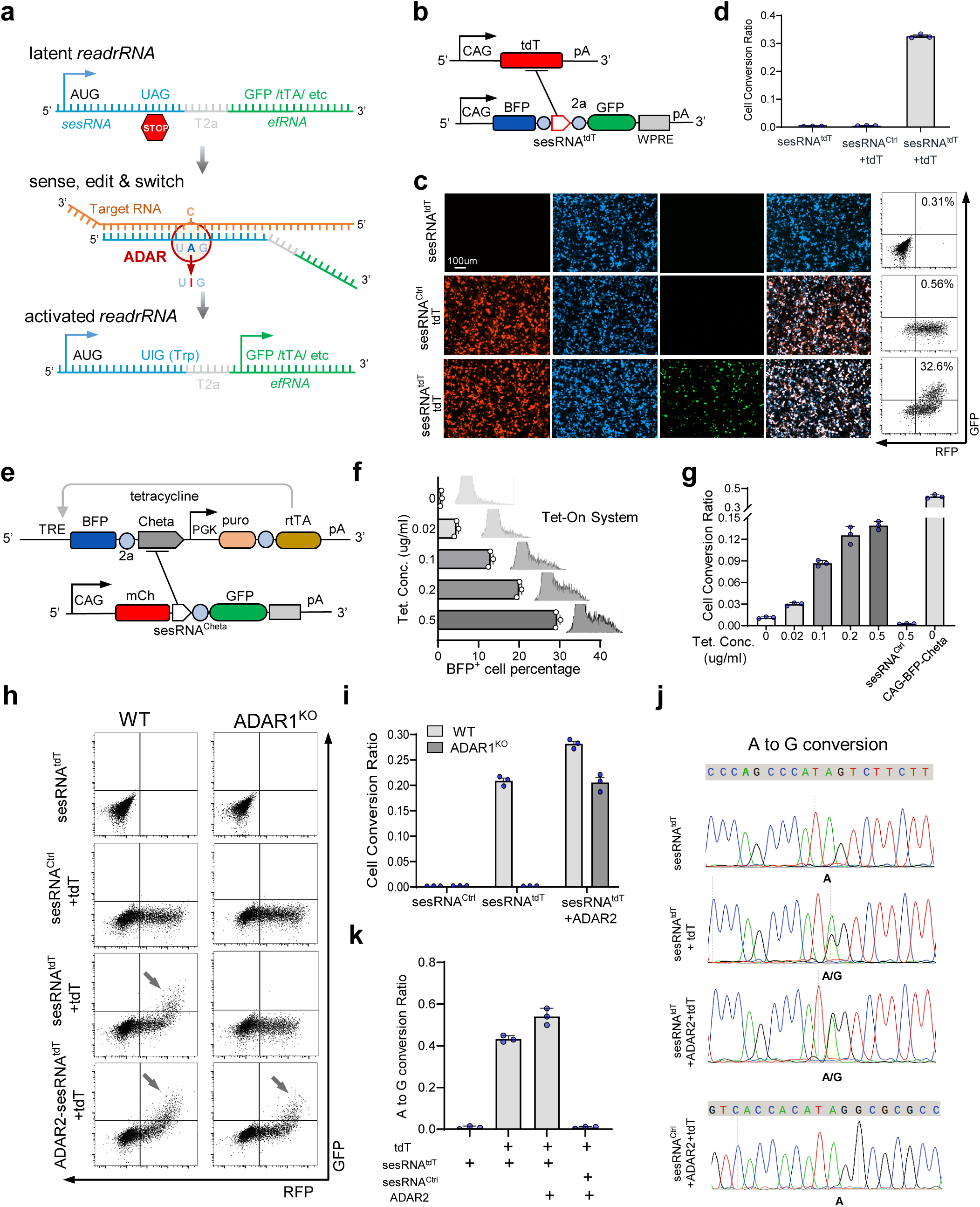
CellREADR design and implementation in mammalian cells. **a,** CellREADR is a single modular readrRNA molecule, consisting of a translationally in-frame 5’-sensor domain (sesRNA) and 3’-effector domain (efRNA), separated by a T2a coding region. sesRNA is complementary to a cellular RNA and contains an in-frame STOP codon that prevents efRNA translation. Base pairing between sesRNA and target RNA recruits ADARs, which mediate A→I editing and convert the UAG STOP to a UGG Trp codon, switching on translation of effector protein. **b-d, b**, *CAG*-*tdT* (top) expresses the *tdTomato* target RNA from a CAG promoter and *READR^tdT-GFP^* (bottom) expresses a readrRNA consisting of a BFP coding region followed by sesRNA^tdT^ and efRNA^GFP^, driven by a CAG promotor. **c**, 293T cells transfected with both *CAG*-*tdT* and *READR^tdT-GFP^* showed robust GFP expression co-localized with BFP and RFP (bottom). Cells transfected with *READR^tdT-GFP^* only (top) or *CAG*-*tdT* and control *READR ^Ctrl^* expressing a sesRNA^Ctrl^ with a scramble coding sequence containing a STOP codon (middle), showed almost no GFP expression. Representative FACS analysis of GFP and RFP expression is shown to the right. **d**, Quantification of CellREADR efficiency by FACS. **e-g,** Effect of target RNA levels on CellREADR. **e**, Expression vectors for quantifying the effect of target RNA levels on CellREADR. In *rtTA-TRE-ChETA* (top), a BFP-ChETA fusion target RNA is transcribed from TRE, driven by constitutively expressed rtTA in a tetracycline concentration dependent manner. *READR^ChETA-GFP^* (bottom) expresses readrRNA consisting of a mCherry coding region followed by sesRNA^ChETA^ and efRNA^GFP^. **f,** In 293T cells co-transfected with *rtTA-TRE-ChETA* and *READR^ChETA-GFP^*, increasing tetracycline concentrations in medium resulted in increased percentage of BFP^+^ in RFP^+^ cells revealed by FACS analysis. **g,** GFP^+^ to RFP ^+^ ratio increased with increasing tetracycline concentration. *CAG-ChETA* constitutively expressing BFP-ChETA RNA served as a positive control, and *READR ^Ctrl^* expressing sesRNA^Ctrl^ served as a negative control. **h-i,** ADAR1 is necessary for CellREDAR function. FACS analysis (**h**) of wild-type or ADAR1^KO^ 293T cells expressing sesRNA^tdT^ only, sesRNA^Ctrl^ and tdT, sesRNA^tdT^ and tdT, or sesRNA^tdT^ and tdT with ADAR2 over expression. Arrows indicate GFP ^+^/RFP^+^ populations. **i**, Quantification of cell conversion ratio. **j,** Electropherograms of Sanger sequencing showing A-to-G conversion at the intended editing site in different samples as indicated. **k**, Bar graph shows the quantification of the A-to-G conversion rate at the targeted editing site. Error bars in **d**, **f**, **g**, and **k** are mean values□±□s.e.m. n□=□3; n represents the number of independent experiments performed in parallel.

We first built a proof-of-principle version of CellREADR and tested it in human 293T cell line. We used an expression vector (*PGK*-*tdT*) to express the *tandem-dimer-Tomato* gene as an exogenous target RNA and to label transfected cells. We then designed a *READR* vector (*READR^tdT-GFP^*) expressing a readrRNA consisting of a 5’ sensor region to the *tdT* sequence embedded with a UAG STOP codon (sesRNA^tdT^), a T2A coding sequence, and a 3’ coding cassette for GFP (**Extended Data Fig. 1a, b**). Co-transfection of *READR^tdT-GFP^* with *PGK*-*tdT* resulted in GFP expression in a subset of tdT*^+^* cells (**Extended Data Fig. 1c**, **d**). To substantiate this result, we replaced the GFP coding region of *READR* vector with that of a tetracycline inducible transcription activator (tTA2), which drove the expression of a firefly luciferase gene (Luc) from the tetracycline response element (*TRE*) promoter (**Extended Data Fig. 1e**). Co-transfection of *READR^tdT-tTA2^* and *TRE-Luc* with *PGK*-*tdT* resulted in approximately 8-fold induction of luminescence compared to empty vector control (**Extended Data Fig. 1f**). We observed sparse and weak GFP or luciferase expression, respectively, from *READR^tdT-GFP^* alone or *READR^tdT-tTA2^*/ *TRE-Luc* when co-transfected with an empty PGK vector (**Extended Data Fig. 1b-f)** which may have resulted from occasional read-through of the STOP codon^28^ in the sesRNA. Inclusion of an in-frame spacer region of approximately 600bp before the sesRNA virtually eliminated residual GFP translation (**Extended Data Fig. 1g, h**); longer spacers decreased the efficacy of readrRNA in the presence of tdT target RNA (**Extended Data Fig. 2h**).

To more quantitatively characterize CellREADR efficiency, we inserted a BFP coding cassette upstream of the sesRNA^tdT^ region, which functioned as a spacer as well as a marker of transfected cells (**Fig. 1b, c**). We co-transfected this *READR^tdT-GFP^* construct with *PGK*-*tdT* in 293T cells, and used FACS for quantitative analysis. We gated on BFP-positive cells to calculate CellREADR efficiency, given that every BFP-positive cell has the potential to express GFP. In the absence of tdT target RNA, *READR^tdT-GFP^* showed almost no GFP expression (0.31%), indicating no detectible STOP read-through. In the presence of *tdT* RNA, the efficiency of *READR^tdT-GFP^* (i.e. the ratio of GFP and RFP double positive cells among RFP positive cells) was 32.6% (**Fig1. c, d**), while a control *READR^Ctrl-GFP^* vector expressing a sesRNA^Ctrl^ with scramble coding sequence showed almost no GFP translation (0.46%). The efficiency reached 67.3% when a TRE/tTA2 system was used for amplification (**Extended Data Fig. 1i-j)**. Next, we examined whether CellREADR efficiency correlated with target RNA levels. The efficiency of *READR^tdT-GFP^* increased with increasing amount of *CAG-tdT* vector in 293T cell transfection (**Extended Data Fig. 3 a, b**). We further designed a tetracycline inducible expression vector in which a ChETA (a variant of light-gated channelrhodopsin-2) coding sequence was fused to BFP and driven by TRE, thus BFP fluorescence level indicated ChETA transcript level, which correlated to tetracycline concentration in culture medium (**Fig. 1e**). Increase of tetracycline concentration and BFP expression levels increased *READR^ChETA-GFP^* efficiency in a ChETA RNA dependent manner (**Fig. 1g** **and Extended Data Fig. 3c, d**). With a constitutive promoter driven BFP-ChETA expression, *READR^ChETA-GFP^* efficiency reached as high as 41.5% (**Fig. 1g** **and Extended Data Fig. 3e, f**). These results demonstrated the target RNA level dependence of CellREADR efficiency.

To demonstrate the capacity of effector RNAs for mediating physiological function, we constructed *READR^tdT-Cas9^* to express the Cas9 protein for genomic DNA editing^29^ and *READR^tdT-taCas3/TEV^* to express functional Caspase3 protein for programmed cell death^30^. Efficient gene editing (**Extended Data Fig. 2a-d)** and increased cell apoptosis (**Extended Data Fig. 2e-f)** was induced, respectively, only with the presence of tdT target RNA.

To examine the role of ADAR proteins in CellREADR function, we generated an *ADAR1* knockout (KO) 293T cell line using CRISPR^31^ (**Extended Data Fig. 4a, b**). As ADAR1 is highly expressed while ADAR2 is barely expressed in 293T cells^25,27,32^, ADAR1^KO^ 293T essentially represented an ADAR-null cell line. Removal of *ADAR1* largely eliminated *READR^tdT-GFP^* functionality (**Fig. 1h, i**), which was rescued by exogenous expression of either the p110 or p150 ADAR1 isoform (**Extended Data Fig. 4c-d**) and the ADAR2 protein (**Fig. 1h, i**). Overexpression of ADAR2 in wild type cells mildly increased *READR^tdT-GFP^* efficiency by ∼25% (**Fig. 1h, i**). Sanger sequencing confirmed A-to-I/G editing at the intended site that converted the TAG STOP codon to a TGG tryptophan codon (**Fig. 1j, k**). Since ADAR expression can be induced by interferon stimulation ^32,33^, we confirmed this induction in 293T cells and further demonstrated that interferon treatment increased *READR^tdT-GFP^* efficiency (**Extended Data Fig. 4e-h**).

In addition to 293T cells, we demonstrated CellREADR functionality in several other cell lines originated from different tissues or species, including human HeLa, mouse N2a, and mouse KPC1242 cell lines (**Extended Data Fig. 5**). Collectively, these results demonstrate that by leveraging cell-endogenous ADARs, CellREADR can couple the detection of cellular RNAs to the translation of effector proteins to manipulate the cell in a way that is specific, efficient, and robust.

### sesRNAs properties confer CellREADR programmability

As sesRNA is a key component of CellREADR, we explored several properties of sesRNAs using the *READR^tdT-GFP^* vector (**Fig. 2a**). We found that sesRNA of less than 50 nucleotides were largely ineffective, while sesRNAs of increasing length positively correlated with the CellREADR efficiency, with an optimal length of 200-350nt (**Fig. 2a**). In terms of sequence mismatch with target RNA, sesRNAs of ∼200nt tolerated up to 10 mismatches (5% of sequence length) or in-frame indels without major decrease of *READR^tdT-GFP^* efficiency (**Fig. 2b** **and Extended Data Fig. 6**); this property confers flexibility for sesRNA design (see **Methods**). However, mismatches near the editing site reduced CellREADR efficiency (**Extended Data Fig. 6**). To explore the effect of locations along target transcript on sesRNAs design, we used an *EF1a-ChETA-tdT* expression vector and tested 5 sesRNAs targeting the promotor and different coding regions. All 4 sesRNAs targeting the transcript region exhibited robust efficiency whereas the sesRNA targeting non-transcribed promoter region did not (**Fig. 2c**). Finally, we examined whether inclusion of more STOP codons would further increase sesRNA stringency (i.e. reduce basal translation in the absence of target RNA, **Fig.1d**). Using luciferase as a sensitive indicator of leaky translation (**Fig. 2d**), sesRNAs with two STOP codons showed reduced leakage with similar efficiency as those with one STOP codon (**Fig. 2e**), although 3 STOP codons significantly reduced the efficiency (**Fig. 2f**). Together, these results suggest strategies to enhance stringency, efficiency and flexibility of CellREADR.

**Fig 2:**
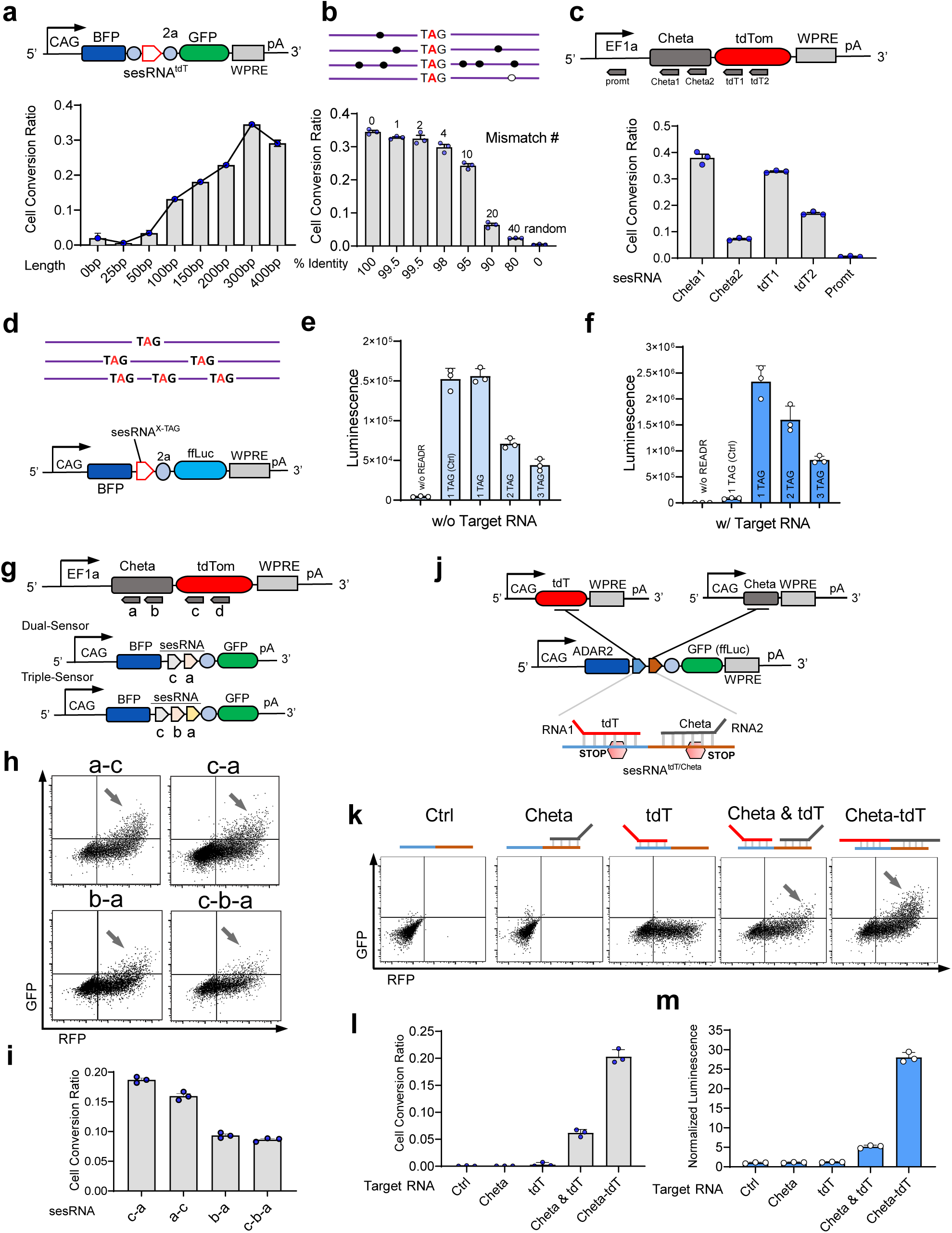
sesRNA properties. **a,** The optimal sesRNA length is ∼200-300 nt. Using *READR^BFP-tdT-GFP^* and tdT as target RNA, sesRNA^tdT^ with variable lengths were tested to identify an optimal length, quantified as cell convertion ratio in FACS assay. **b,** Effect of mismatches between sesRNA and target RNA base pairing. Number of mismatches (top) and percent identity between sesRNA^tdT^ and tdT target RNA (bottom) are shown for each case. **c,** Effect of sesRNA sensing of different locations and sequences within the target RNA (e.g. different ChETA and tdT coding regions) expressed from a *EF1a-ChETA-tdT* vector; promoter region was used as negative control. readrRNA efficiencies were shown (bottom). **d,** To improve stringency, sesRNA^tdT^ was designed to contain 1 to 3 STOP codons (X-TAG) in *READR^tdT-Luc^* vector. **e**-**f,** Luminescence of 293T cells transfected with *READR^tdT-Luc^* only (**e**) or co-transfected with *CAG-tdT* (**f**). **g,** Schematic of dual and triple sensor *READR^ChETA/tdT-GFP^* vectors with sesRNA arrays targeting different regions of a ChETA-tdT fusion transcript expressed from *EF1a-ChETA-tdT.* **h,** FACS analysis of 293T cells co-transfected *EF1a-ChETA-tdT* and a dual or triple *READR^tdT-GFP^* vector. Arrows indicate GFP ^+^/RFP ^+^ populations. **i,** Quantification of the efficiency of various dual or triple *READR^tdT-GFP^* vector in (**h**). **j,** Scheme for intersectional targeting of cells expressing two target RNAs (ChETA and tdT) using a dual-sensor *READR^ChETA/tdT-GFP^* or *READR^ChETA/tdT-Luc^* vector; each sesRNA in the dual-sensor array contains an editable STOP. **k,** FACS analysis showed that co-transfection of 293T cells with *READR^Chet/atdT-GFP^* and either *CAG-ChETA* or *CAG-tdT* resulted in almost no GFP translation, and only triple transfection resulted in GFP expression. *EF1a-ChETA-tdT* (expressing a ChETA-tdT fusion transcript) co-transfection was used as a positive control. Quantifications were shown with FACS (**l**) and luciferase assay **(m)**. Error bars in **a-c**, **e-f**, **i**, and **l-m** are mean values□±□s.e.m. n□=□3, n represents the number of independent experiments performed in parallel.

Built on Watson-Crick base-pairing, CellREADR is inherently programmable, conferring potential for joint sensing of two or more different RNAs to access more specific cell types defined by RNA co-expression. To explore this property, we first designed dual or triple sesRNA arrays, with each sesRNA containing an editable STOP codon, that target different regions of the same transcript (**Fig. 2g, j**). All of the tested sesRNA arrays exhibited robust CellREADR efficiency, comparable to that of a single sesRNA (**Fig. 2h, i**). To examine the possibility for joint detection of two target RNAs, we designed a sesRNA array consisting of tandemly arranged sesRNA^tdT^ and sesRNA^ChETA^ (sesRNA^tdT/ChETA^) (**Fig. 2j**). Whereas single target RNA alone, either tdTomato or ChETA, failed to induce GFP or Luc expression with readrRNA^tdT/ChETA^, significant GFP expression or increased luminescence were observed, respectively, when both tdTomato and ChETA vectors were transfected (**Fig. 2k-m**). This result demonstrated the potential for intersectional targeting of more specific cell types based on the programmability of two or more RNA sensors in CellREADR.

### Endogenous RNA sensing by CellREADR

Compared with synthetic genes expressing exogenous RNAs, endogenous genes often have more complex genomic structures, including numerous exons, introns, and regulatory elements; transcribed endogenous RNAs undergo multiple and elaborate post-transcriptional steps such as splicing and chemical modifications before being processed as mature mRNAs^34^. To examine the capacity of CellREADR for detecting cell endogenous RNAs, we first selected *EEF1A1*, a housekeeping gene highly expressed in 293T cells, and systematically designed a set of sesRNAs targeting its various exons, introns, 5’ and 3’ UTR, and mRNAs (**Fig. 3a, b**). These sesRNAs were effective in sensing all intended *EEF1A1* regions to switch on efRNA translation; exons appeared to be better targets than introns (**Fig. 3c**). Notably, sesRNAs complementary to a mRNA region joined from two spliced exons achieved comparable efficiency as to those contained within an exon, indicating that sesRNA can be designed to target both pre-mRNA and mRNA sequence. CellREADR efficacy to *EEF1A1* RNAs increased with incubation time after cell transfection (**Extended Data Fig. 7**).

**Fig 3.**
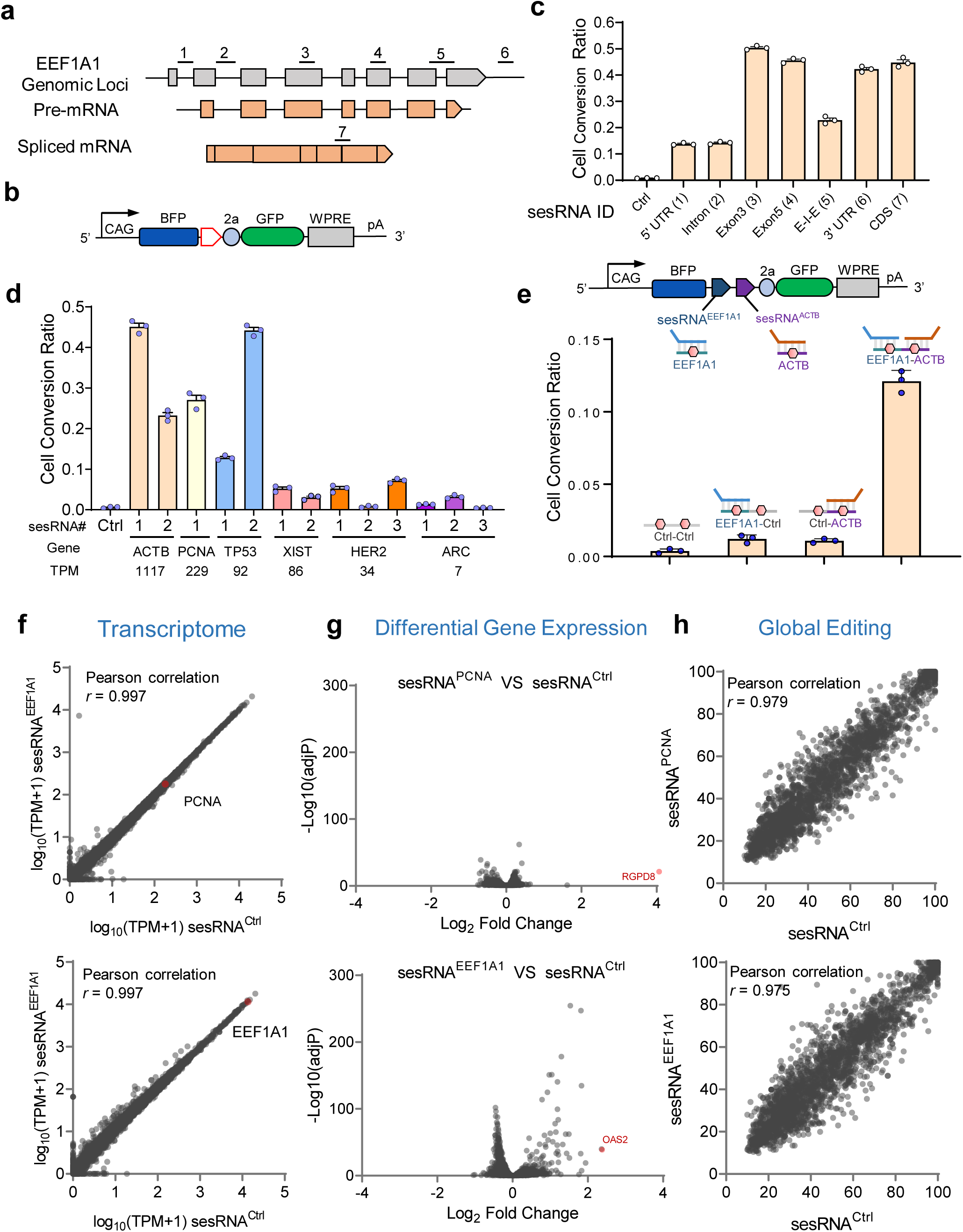
Endogenous RNA sensing with CellREADR. **a,** Schematic of the genomic loci of human *EEF1A1* gene, showing exon and intron structures (not in scale), pre-mRNA (middle), and mRNA (bottom). sesRNA^EEF1A1^ (1-7) were designed to sample across the *EEF1A1* transcripts including the coding sequence (CDS). **b,** Schematic of the *READR^EEF1A1-GFP^* vector for testing sesRNA^EEF1A1^ (1-7); the *BFP* expression cassette labels all transfected cells. **c,** Quantification of the efficiency of sesRNA^EEF1A1^ (1-7) in 293T cells. **d,** High sensitivity of CellREADR. Quantification of CellREADR efficiency for several endogenous cellular RNAs with different expressing levels; TPM (transcript per million) from RNAseq data was used to indicate cellular RNA expression levels. 1-3 sesRNAs were designed for each target. **e,** Schematic for intersectional targeting of two endogenous RNAs (EEF1A1 and ACTB) using a dual-sensor *READR^EEF1A1/ACTB-GFP^* vector; each sesRNA in the dual-sensor array contains an editable STOP. Dual-sensor *READR^Ctr1/Ctrl-GFP^, READR^EEF1A1/Ctrl-GFP^* or *READR^Ctr1/ACTB-GFP^* were used as controls. Quantifications of efficiency showed that only 293T cells with *READR^Chet/atdT-GFP^* transfection resulted in significant GFP expression. **f,** CellREADR did not alter cellular transcriptome assayed by RNA sequencing. Comparisons of transcriptomes between cells transfected with sesRNA^EEF1A1(7)^ or sesRNA^Ctrl^ (top) and with sesRNA^PCNA^ or sesRNA^Ctrl^ (bottom). **g.** Volcano plot of differential gene expression analysis between sesRNA^EEF1A1^ or sesRNA^Ctrl^ (top), sesRNA^PCNA^ or sesRNA^Ctrl^ (bottom). Genes with adjusted P value < 0.01 and log_2_ (fold change) > 2 were defined as significantly differentially expressed genes, labelled in red. One gene OAS2 was detected with significantly increased expression in the sesRNA^EEF1A^ group. **h,** Transcriptome-wide analysis of the effects of sesRNA^EEF1A1^ (top) or sesRNA^ACTB^ (bottom) on A-to-I editing by RNA-sequencing. Pearson′s correlation coefficient analysis was used to evaluate the differential RNA editing rate. Error bars in c-e are mean values□±□s.e.m. n□=□3, n represents the number of independent experiments performed in parallel.

To examine the sensitivity of CellREADR to target RNA levels, we selected several endogenous RNAs with expression ranging from high (*ACTB, PCNA*), modest (*TP53, XIST*), to low (*HER2, ARC*) levels. CellREADR was able to reliably detect even low level *ARC* RNAs, although at much lower efficiency compared to modest or high level RNAs (**Fig. 3d**). For the same RNA expressed at different levels, sesRNA sequence was a major determinant of RNA sensing (**Fig. 3d**).

To explore intersectional detection of two endogenous RNAs, we designed a sesRNA array consisting of tandemly arranged sesRNA^EEF1A1^ and sesRNA^ACTB^. sesRNA^EEF1A1/ACTB^ triggered robust effector GFP translation comparing to control sesRNA arrays that included one or two scramble sesRNAs (sesRNA^EEF1A1/Ctrl^, sesRNA^Ctrl/ACTB^, sesRNA^Ctrl/Ctrl^; **Fig. 3e**). This result strengthens the potential of CellREADR for more specific targeting of cell types by intersectional RNA sensing.

To evaluate the effect of CellREADR on cellular gene expression, we used RNA-sequencing to compare the transcriptome profiles of 293T cells expressing sesRNA^EEF1A^ or sesRNA^PCNA^ with those expressing a scrambled sesRNA^Ctrl^ . Correlation analysis^35^ showed that sesRNA^EEF1A^ or sesRNA^PCNA^ expression had no significant impact on global transcriptome (**Fig. 3f**). Differential expression analysis^36^ revealed that sesRNA^PCNA^ led to a reduction of only one gene with no clear link to *PCNA* (e.g. *RGPD8*). sesRNA^EEF1A1^ resulted in a ∼4-fold increase of OAS2 mRNA (2’-5’-Oligoadenylate Synthetase 2, which encodes an interferon-induced and dsRNA-activated antiviral enzyme, **Fig. 3g**) and more mild increase (2 to 4-fold) of several other immune-related mRNAs (**Extended Data Table 3)**. These results suggest that sesRNAs, even those targeting high level transcript (EEF1A1), exert no significant activation of cellular innate immune response, unlike cells transfected with synthetic analog of dsRNA-poly(I:C)^25^.

To assess whether CellREADR perturbs RNA editing in cellular transcriptomes, we used REDItool^37^ to compare the global A-to-I editing sites between sesRNA^EEF1A1^ or sesRNA^PCNA^ and sesRNA^Ctrl^ groups. Neither the sesRNA^PCNA^ nor sesRNA^EEF1A1^ group showed a significant difference compared to the sesRNA^Ctrl^ group (**Fig. 3h**), demonstrating that CellREADR had no effects on normal A-to-I editing of cellular transcriptomes. Furthermore, CellREADR did not alter the levels of target RNA transcripts assayed by quantitative PCR (qPCR) (**Extended Data Fig. 8a**); this result also ruled out the possible effect of RNA interference induced by readrRNAs. To evaluate the possibility of unintended editing on target RNA, we examined all adenosine sites within the EEF1A1 RNA targeted by sesRNA^EEF1A1^. Very low editing was observed throughout the targeted sequence window (**Extended Data Fig. 8b**). Two adenosines around the A-C mismatch site (intended editable STOP in sesRNA) showed higher editing rate, but at less than 1% **(Extended Data Fig. 8c)**. Together these results indicate that cell endogenous ADARs enable efficient and robust CellREADR functionality with minimum effects on cellular gene expression, RNA editing, and target transcript.

### Neuronal cell type targeting with CellREADR

To apply CellREADR for accessing specific cell types in animal tissues, we designed both a singular and a binary vector system (**Fig. 4b**). In singular READR vector, a hSyn promoter drives transcription of mCherry followed by sesRNA and efRNA coding for smFlag and tTA2. This vector directly couples RNA detection with tool gene expression; the efficiency and specificity can be evaluated by co-localization of mCherry, smFlag, and the target mRNA or protein. The binary system includes an addition Reporter vector, which contains a TRE promoter driving mNeonGreen (mNeon) or other effector proteins in response to tTA2 translated from the *READR* vector (**Fig. 4b**). The binary vectors provide amplification of tool gene expression as well as the combinatorial flexibility of expressing different tool genes in different cell types by pairing *READR* and *Reporter* vectors.

**Fig 4.**
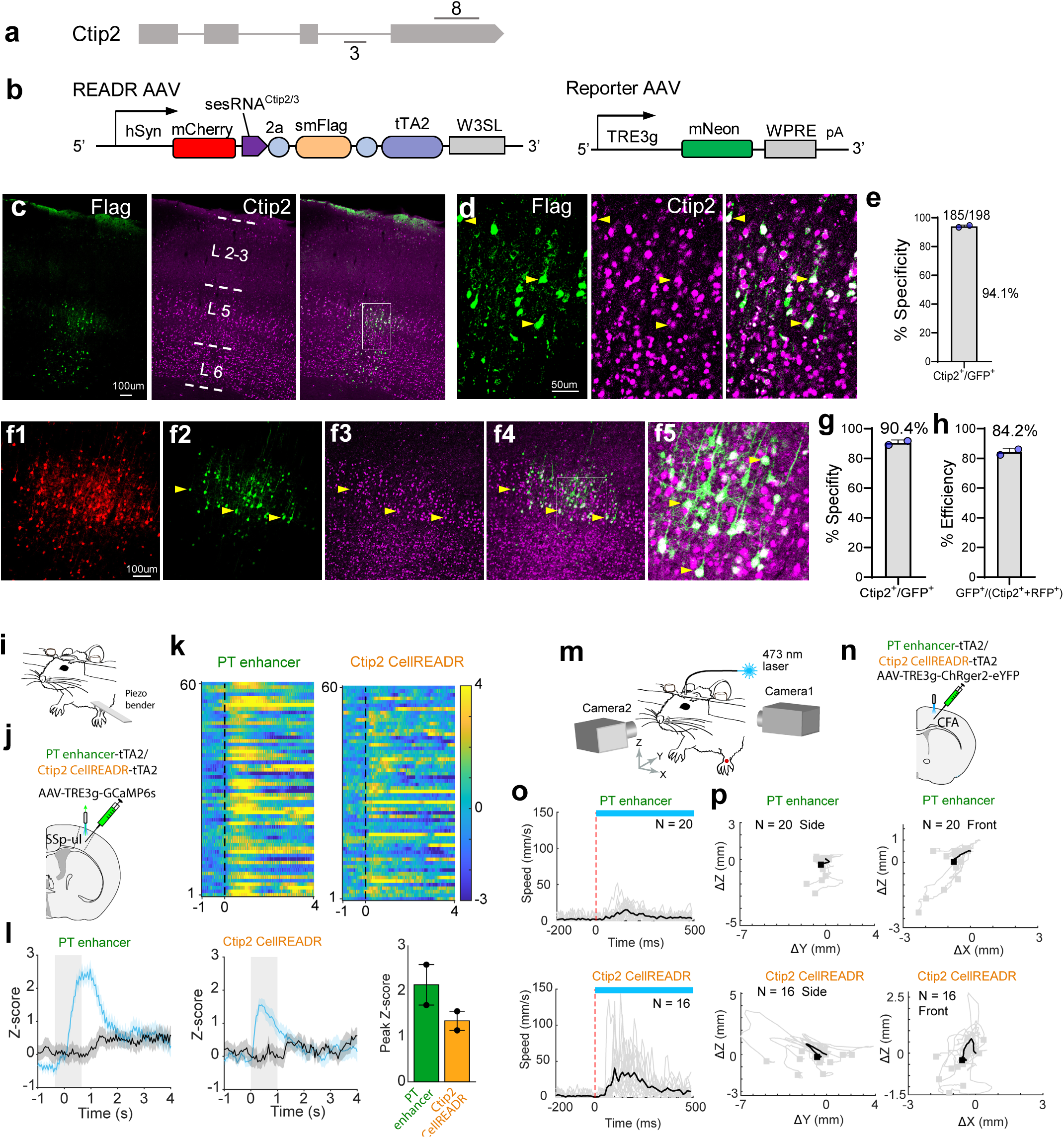
CellREADR targeting, monitoring, and manipulation of a neuronal cell type in mice. **a.** Genomic structure of the mouse *Ctip2* genes with locations of sesRNA3 and sesRNA8 as indicated. **b,** Schematic of singular and binary AAV vectors for targeting *Ctip2* neurons in brain tissues. In singular *READR^Ctip2-smFlag/tTA2^* vector, a hSyn promoter drives expression of mCherry followed by sequences coding for sesRNA^Ctip2^, smFlag and tTA2 effectors. In binary vectors, a Reporter vector drives mNeonGreen expression from a TRE promoter in response to tTA2 from *READR^Ctip2-smFlag/tTA2^*. **c,** Coronal section of S1 cortex injected with *READR^Ctip2(3)-smFlag/tTA2^* AAVs. Immunofluorescence with FLAG (left) and CTIP2 (middle) antibodies indicated that READR-labeled cells (FLAG*^+^*, green) showed high colocalization with CTIP2*^+^* PNs. **d,** Magnified view of boxed region in (**c**). Arrowheads show representative co-labeled cells. **e,** Specificity of *READR^Ctip2(3)^* delivered with singular vector. **f,** In S1 cortex co-injected with binary AAV vectors, neurons infected by *READR^Ctip2^* expressed mCherry (**f1**). In a subset of these neurons, sesRNA^Ctip2(3)^ triggered translation of tTA2 and activation of mNeon expression in L5b PNs (**f**2). The specificity of mNeon expression in Ctip2 PNs was assessed with CTIP2 immunofluorescence (**f3-5**). **f**5 is a boxed region in **f**4. Arrowheads indicate the co-labeled cells. **g,** Specificity of *READR^Ctip2(3)^* delivered with binary vectors. **h,** Efficiency of binary *READR^Ctip2(3)^*, calculated as GFP^+^ cells among mCherry and CTIP2 expressing cells. **i-l,** *READR^Ctip2(3)^* enabled calcium imaging of a specific cortical neuron type in vivo. **i-j,** Schematic of left-paw stimulation in anesthetized mice (**i**) and fiber photometry recording (**j**). **j,** AAV-PTenhancer-tTA2 or AAV-Ctip2-CellREADR-tTA2 was co-injected with AAV-TRE-GCaMP6s into primary somatosensory cortex, upper limb area (SSp-ul). **k,** Heatmap of neuronal activity aligned to the onset of left-paw stimulation. **l,** Mean calcium signal aligned to the onset of left-paw stimulation (left two panels). Peak Z-score of the responses was shown (right panel, n = 2 mice each). Gray area indicates 1s of stimulation. Black traces are from data aligned to shuffled onset times. Shades around mean denote ± s.e.m. Data are mean ± s.e.m. **m-p,** *READR^Ctip2(3)^* enabled optogenetic manipulation of a specific cortical neuron type in behaving mice. **m-n,** Schematic of optogenetic stimulation experiment in head-fixed mice (**m**) and viral injection (**n**). Two cameras were used to record induced movements. Nose tip is the coordinate origin. X, Y, and Z axes correspond to medial-lateral, anterior-posterior, and dorsal-ventral axes, respectively. A reflective marker was attached to the back of left paw for tracking (red dot). **n,** AAV-PTenhancer-tTA2 or AAV-Ctip2-CellREADR-tTA2 was co-injected with AAV-TRE-ChRger2-eYFP into caudal forelimb area (CFA). **o,** Speed of left paw before and after stimulation onset (n = 20 trials for PT enhancer mouse; n=16 trials for Ctip2 CellREADR mouse). Black traces indicate the average. Red line denotes stimulation onset. Blue bar represents 500 ms of stimulation (also **see suppl video 1**). **p,** Movement trajectories of left paw during optogenetic stimulation (n = 20 trials for PT enhancer mouse; n = 16 trials for Ctip2 CellREADR mouse). Side (left) and front (right) trajectories were normalized to the start position of left paw. Black trajectories indicate the average. Squares indicate left-paw positions when stimulation stopped.

We first designed a set of *READR* vectors targeting mRNAs that define major glutamatergic projection neuron (PN) types of the mouse cerebral cortex. The zinc-finger transcription factor *Fezf2* labels the vast majority of layer 5b (L5b) and L6 corticofugal projection neurons (CFPNs) that constitute cortical output channels, while *Ctip2* predominantly labels a subset of L5b/L6 CFPNs neurons^38,39^. To identify appropriate sesRNAs, we first carried out a screen in 293T cells by co-expressing individual sesRNAs with their target sequences (**Extended Data Fig. 9a, b, d**). We designed 4 sesRNAs targeting exon1 or 5’UTR of *Fezf2* (**Extended Data Fig. 9a,c**) and 8 sesRNAs targeting various exons and introns of *Ctip2* (**Extended Data Fig. 9b,c**). All sesRNAs exhibited variable but significant CellREADR efficiency in 293T cells in the presence of exogenous *Fezf2* or *Ctip2* target sequences (**Extended Data Fig. 9e-f**).

We then used AAV vectors to test these sesRNAs in mouse cortex by focal injection into the whisker barrel somatosensory cortex (S1) or motor cortex (M1). Delivered with binary vectors, *Fezf2* sesRNA1 showed 86.4% specificity, and *Ctip2* sesRNA3 and 8 reached to over 90% specificity assayed by *Ctip2* antibody staining. (**Extended Data Fig. 9j-k**). We then focused on further evaluating *Ctip2* sesRNA3. In S1 cortex infected with singular *READR^Ctip2/3^*AAV, FLAG immunofluorescence was concentrated in deep layers (**Fig. 4c**); co-localization of CTIP2 and FLAG immune-labeled cells (**Fig. 4c, d**) showed a specificity of 94.1% (**Fig. 4e**). In S1 cortex co-infected with binary *READR^Ctip2/3^*/*Reporter ^mNeon^* AAVs, mCherry expression from the READR vector labeled cells across cortical layers, while mNeon expression from the Reporter vector specifically labeled L5b and some L6 neurons (**Fig. 4f**); CTIP2 immunofluorescence showed a specificity of 90.4% (**Fig. 4g**). The efficiency of *READR^Ctip2/3^*, calculated as the ratio of GFP cells among mCherry and Ctip2 double positive cells, reached 84.2% (**Fig. 4h**).

To test whether ADAR2 overexpression from the READR vector might enhance CellREADR functionality, we replaced mCherry in *READR* vector with an *ADAR2* cDNA, and tested sesRNAs for *Fezf2* and *Ctip2* in mouse cortex (**Extended Data Fig. 10a-c**). *ADAR2* overexpression did not notably enhance the specificity or efficiency compared with binary vectors that relied on endogenous *ADAR* (**Extended Data Fig. 10c-i**).

In addition to *Ctip2* and *Fezf2* mRNAs that mark CFPNs, we designed sesRNAs in binary vectors to target several mRNAs expressed at a range of levels^40^ (**Extended Data Fig. 11**), including *PlxnD1* (intratelencephalic-IT PNs in L2/3 and L5a), *Satb2* (IT PNs across both upper and lower layers), *Rorb* (L4 pyramidal neurons), and vGAT (pan-GABAergic neurons), respectively. Although only 2-3 sesRNA were tested in each case, we were able to identify sesRNAs that achieved at or above 75% specificity (**Extended Data Fig. 12**). These results indicate that CellREADR is a robust method, and specific cell type targeting can be achieved by testing multiple sesRNAs for each target.

To evaluate the long-term effects of CellREADR sesRNAs in vivo, we incubated *READR^Ctip2/3^* and a control *CAG-tdT* AAVs in mouse cortex for 3 months. Quantitative RT-PCR assay on expression of nine genes implicated in glia activation and immunogenicity showed no significant change in *READR^Ctip2/3^*-infected versus control tissues (**Extended Data Fig. 13**).

### Recording and control of neuron type with CellREADR

In addition to specificity, a key feature of any cell type technology is its efficacy for monitoring and manipulating cell function. We tested CellREADR efficacy by applying it to record and control neuronal function in mice. As a benchmark, we compared CellREADR with transcription enhancer-based cell type targeting approach. The mscRE4 enhancer was identified, through a large scale open-chromatin screening and in vivo validation effort, that labels L5 pyramidal track (PT) neurons, a subset of CFPNs^18^. We thus compared the efficacy of mscRE4 enhancer (or PT enhancer) with Ctip2 sesRNA3 by using the same tTA-TRE binary vectors for expressing genetic-encoded calcium indicator GCaMP6s and light-activated ion channel channelrhodopsin2 (**Fig. 4i-p**).

In mice co-infected with *PT enhancer-tTA2*/*Reporter^GCaMP6s^* or *READR^Ctip2/3^*/*Reporter^GCaMP6s^* AAVs in the forelimb somatosensory cortex, mechanical stimulation of the forepaw induced reliable and time-locked increase of GCaMP6s signals in forelimb somatosensory cortex, measured by fiber photometry, in both PT enhancer and *READR^Ctip2/3^* infected mice (**Fig. 4i-l**). This result indicates that tool gene expression levels achieved by CellREADR were sufficient to monitor cell physiology in live animals.

In mice co-infected with *PT enhancer-tTA2/Reporter^ChRger2-eYFP^* or *READR^Ctip2/3^/Reporter^ChRger2-eYFP^* AAVs in the caudal forelimb motor area (CFA), light stimulation of CFA induced reliable and time-locked forelimb movement, which appeared stronger in *READR^Ctip2/3^* compared to PT enhancer infected mice (**Fig. 4m-p, Extended Data Movie. 1**). This result indicates that tool gene expression levels achieved by CellREADR were sufficient to control cell function in behaving animals. Anatomical analysis of eYFP expression confirmed L5 PT neuron labeling by *READR^Ctip2/3^*, as revealed by axonal projections in subcortical targets including striatum, thalamus, pons, and medulla (**Extended Data Fig. 14**).

### CellREADR targeting of neuron types across mammalian species

To demonstrate the generalizability of CellREADR across mammalian species, we performed a series of experiments in rat and human brain. We first targeted GABAergic neurons in rats by using binary AAV vectors targeting vGAT mRNA, a pan-GABAergic transcript (**Extended Data Fig. 15b**). Co-injection of *READR^vGAT^* (*hSyn-mCherry -sesRNA^vGAT^-smFlag-tTA2)* with *Reporter^mNeon^* (*TRE3g-mNeon)* AAVs (**Extended Data Fig. 15a**) into rat S1 barrel cortex and hippocampus specifically labeled GABAergic interneurons in the cortex (**Extended Data Fig. 15c**) and hippocampus (**Extended Data Fig. 15d-c)**. The specificity, assayed by co-labeling of *vGAT* mRNA (magenta) and *READR^vGAT^* (GFP), was 91.7% for cortex and 93.8% for hippocampus, respectively **(Extended Data Fig. 15f-g)**. Tle4 is a conserved transcription factor that marks L6 corticothalamic (CT) pyramidal cells in rodents. A sesRNA targeting exon15 of *Tle4* mRNA (**Extended Data Fig. 15h)** delivered by binary *READR^Tle4^* AAVs labeled deep layer neurons with a co-labeling by TLE4 antibody at 62.5% (**Extended Data Fig. 15i-k)**.

We additionally established an oragnotypic culture platform for studying CellREADR in human neocortical specimens collected from epilepsy surgeries^41^ (**Extended Data Fig. 16a)**. First, using a GFP construct driven by the hSyn promoter (*AAVrg-hSyn-eGFP*), we observed dense viral labeling across multiple layers, with a multitude of observed morphologies and a predominance of pyramidal neurons characterized by prominent apical dendrites (**Extended Data Fig. 16b, c**). We then designed two sesRNAs targeting *FOXP2* mRNA (Forkhead box protein P2) (**Fig. 5a, b**), an evolutionarily conserved gene expressed in cortical and striatal projection neurons and implicated in human language skill development^42^. In situ hybridization and immunostaining indicate that, unlike in mouse, where *FoxP2* expression is restricted to L6 CT neurons^39^, human FOXP2 is expressed across cortical layers (**Extended Data Fig. 16d**). Five days after applying binary *READR^FOXP2^/Reporter^mNeon^* AAVs (*hSyn-ClipF–sesRNA^FOXP2^-smV5-tTA2* with *TRE3g-mNeon*) to human neocortical slices, we observed mNeon-labeled cell bodies in upper and deep layers (**Fig. 5c, Extended Data Fig. 16e)**. FOXP2 immunostaining showed the specificity of binary *READR^FOXP2^* was nearly 80% (**Fig. 5d, e**). Labeled neurons also exhibited electrophysiological (**Fig. 5f**) and morphological (**Fig. 5g**) properties consistent with glutamatergic pyramidal neurons^43^. Using two singular *READR^FOXP2^* vectors, each with a different sesRNA, we additionally observed over 97% specificity of targeting FOXP2 neurons (**Extended Data Fig. 16f-i)**.

**Fig 5.**
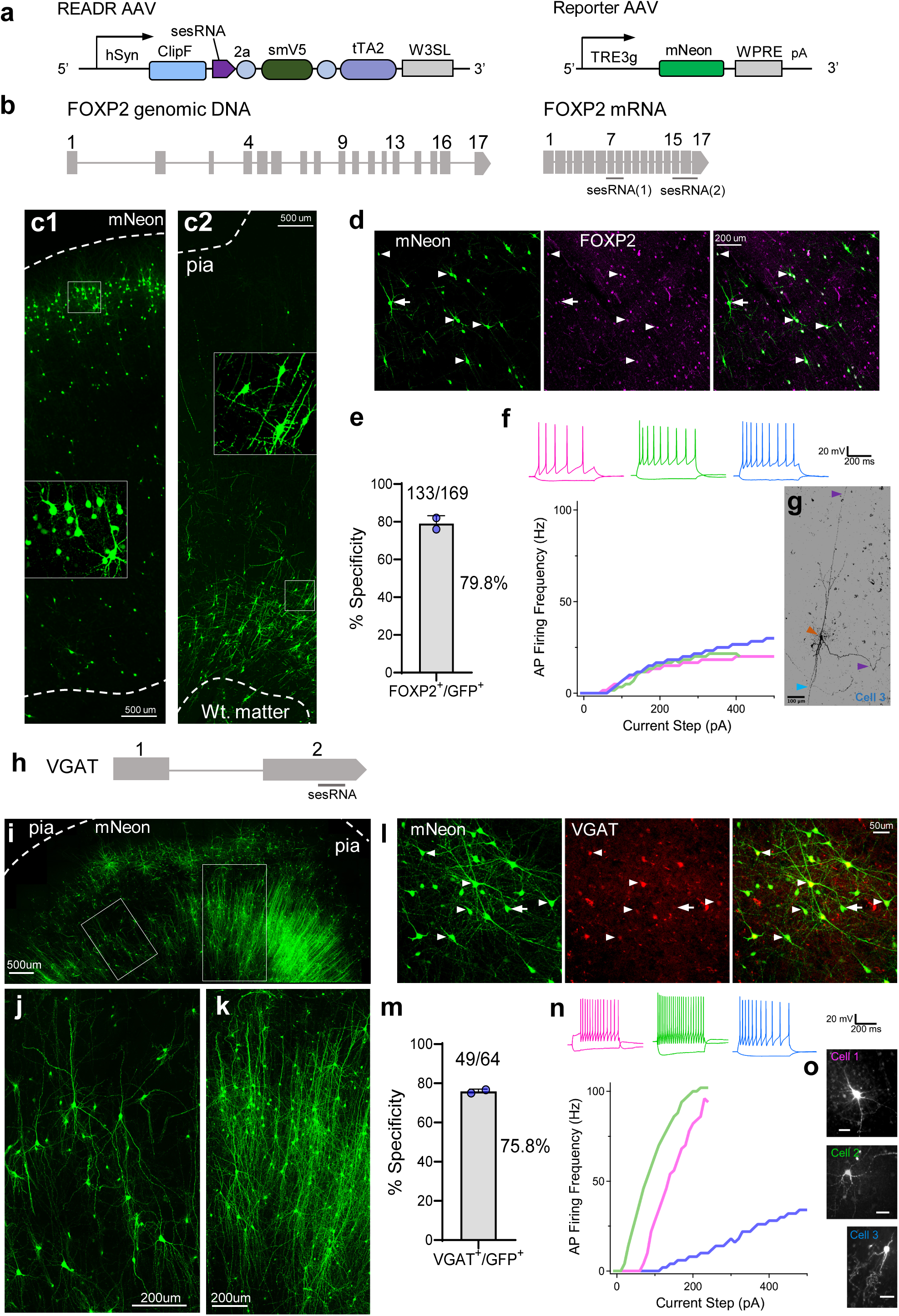
CellREADR-enabled targeting and recording of human cortical neuron types. **a,** Schematic of binary AAV vectors for targeting neuron types in human brain tissues. In the *READR* vector, a hSyn promoter drives transcription of ClipF-Tag followed by sequences coding for sesRNA^FOXP2^, smV5, and tTA2. In *TRE3g-mNeon*, the TRE promoter drives mNeon in response to tTA from the READR virus. **b,** The genomic structure of human FOXP2 gene. Two sesRNAs were designed to target the FOXP2 mRNA. **c,** *READR* targeting of human FOXP2 cells. Upper layer FOXP2 cells (mNeon, native fluorescence) targeted with *READR^FOXP2(1)^* AAVs in organotypic slices from temporal neocortex (**c1**). Deeper layer FOXP2 cells targeted with *READR^FOXP2(2)^* from parietal neocortex (**c2**). Inserts in **c1** and **c2** showed magnified view of boxed region. Dashed lines delineate pia and white matter. **d,** Immunostaining of FOXP2 in *READR^FOXP2(2)^* labeled neurons. Arrowheads indicate neurons co-labeled with mNeon native fluorescence (left) and FOXP2 immunofluorescence (middle); the arrow indicates a cell labeled only by mNeon. **e,** Specificity of *READR^FOXP2(2)^* assayed by FOXP2 immunofluorescence. Error bars are mean values□±□s.e.m. n = 2 biological replicates performed. **f,** Current-clamp recording traces of three mNeon-labeled neurons (top). Input-output curves for cells depict similar spiking behaviors (top). **g,** Morphology of labeled neuron recorded in (**f**) (blue curve). The patched cell was filled with biocytin for post-hoc recovery and was visualized by silver stain in bright field. Brown arrowhead, cyan arrowhead and purple arrowheads indicate the cell soma, axons and dendrites, respectively. **h,** SesRNA designed to target exon2 of the human VGAT mRNA. **i,** An organotypic slice from human temporal neocortex co-infected with AAV*-READR^VGAT^* ;*TRE3g-mNeon* and visualized by native fluorescence 8 days post-infection. **j-k,** Morphologies of AAV*-READR^VGAT^*- labeled interneurons. **l,** Colocalization of mNeon native fluoescence (left) with VGAT mRNA by *in situ* hybridization (middle). Arrowheads indicate co-labeled neurons; arrow indicates a cell labeled only by mNeon. **m,** Specificity of *READR^VGAT^* assayed by VGAT mRNA in situ. Error bars are mean values□±□s.e.m. n = 2 biological replicates performed. **n,** Current-clamp recording traces (top) of three labeled neurons, filled with biocytin for post-hoc recovery. Streptavidin dye was used for visualization. Input-output curves (bottom) depict distinct spiking behaviors (accommodating (blue), fast (green), delayed onset (red)). **o,** Morphologies of *READR^VGAT^* labeled cells recorded in (**n**).

We also targeted human neocortical GABAergic cells by designing a sesRNA to *VGAT* (**Fig. 5h**). Using a binary *READR^VGAT^/Reporter^mNeon^*, we observed cellular labeling across neocortical layers (**Fig. 5i**), with labeled cells exhibiting diverse morphologies characteristic of interneurons, including multipolar and smooth dendrites (**Fig. 5j**), and strikingly dense, vertically oriented axons (**Fig. 5i, k**). *In situ* hybridization for VGAT mRNA showed a specificity of 76.5% (**Fig. 5l, m**), likely an underestimate due to the potentially reduced sensitivity of *in situ* hybridization in cultured *ex vivo* tissues. In support of this possibility, we observed very few labeled cells with pyramidal morphologies (**Fig. 5i-l**). Targeted patch-clamp recordings of *READR^VGAT^/Reporter^mNeon^* labeled interneurons seven days after virus application revealed various intrinsic properties, including accommodating, fast, and delayed onset firing (**Fig. 5n**). The physiological properties (**Fig. 5n**) and partially reconstructed morphologies of the patched cells (**Fig. 5o**) were consistent with those of mammalian neocortical interneurons^44^. Together, these results demonstrate the cross-species generalizability and utility of CellREADR.

## DISCUSSION

The ability to use sequence-programmable RNA sensors as protein translation switches to analyze and manipulate cells define a new generation of molecular and cell engineering tools that will synergize with and expand much beyond enhancer-based methods^13–19^. Based on Watson-Crick base-pairing and endogenous ADAR-mediated RNA editing, CellREADR is inherently 1) specific to cells defined by RNA expression; 2) easy to build, use, and disseminate; 3) scalable for targeting all RNA-defined cells in any tissue; 4) generalizable to most animal species including human; and 5) programmable to achieve intersectional targeting of cells defined by two or more RNAs as well as multiplexed targeting and manipulation of several cell types in the same tissue. Several aspects of the CellREADR can be improved to increase functionality and versatility. For example, development of computational algorithms ^45^ for sesRNA design and their experimental validation should optimize the length, sequence, and STOP codon position for increased specificity and reliability. Improved readrRNA stability and efRNA translation, e.g. by using circular ^46,47^ or chemical modified READR RNAs^24,48^, should boost its efficacy. Innovation in nucleic acid delivery, such as liposome nanoparticles^49^, SEND (selective endogenous encapsidation for cellular delivery) system^29^, and viral capsid proteins that steer tissue and species tropism^50^, will enhance the ease and range of CellREADR applications. Finally, we expect CellREADR to apply to other vertebrate and invertebrate species.

As RNAs are the universal messengers of gene expression that underlie cell identities and cell states in biological processes across life span and species, CellREADR will enable cell type based comparative and evolutionary approaches for discovering general biological principles; it will also accelerate the rapid development of disease models with cell-type resolution, especially using human cells and tissues, for understanding etiology and mechanisms. With the capacity and flexibility to program cellular physiology, CellREADR will facilitate next-generation precision diagnostics and therapeutics, such as detecting cancer cells by RNA markers and eliminating them by inducing programmed cell death or recruiting immune killer cells^51^. While gene editing-^42^ and RNA editing-^52^ based therapeutic approaches have focused almost exclusively on correcting monogenic disease mutations^53,54^, many complex disorders, such as neuropsychiatric and developmental disorders, result from multigenic predispositions that impact the functional wiring of brain circuits and may not be corrected by gene or transcript editing. Modifying and tuning the affected cell types and circuits are rational alternatives^55^. Synergizing with genome and transcriptome engineering technologies, CellREADR will facilitate elucidating the principles of biological information flow from genotype to phenotype through cell types, and usher in a new generation of programmable cell-specific RNA medicine and biotechnology.

## ACKNOWLEDGMENTS

We are grateful to Dr. Ledong Wan for providing the Hela cell line. We thank Drs. Michael Tadross, Anne West, Kate Meyer, Anthony Zador and Scott Soderling for comments on the manuscript. We thank Joshua Hatfield and Bao-xia Han for animal preparation, Raditya Utama for bioinformatic analysis. This work was supported in part by NIH grants 1DP1MH129954-01 and 5U19MH114821-03 to Z.J.H. D.G.S. was supported by the NINDS K12 Neurosurgery Research Career Development Program K12 Award and the Klingenstein-Simons Foundation.

## AUTHOR CONTRIBUTIONS

Y.Q. and Z. J. H. conceived this study. Z.J.H. designed and supervised the research, analyzed data, and wrote the manuscript. Y.Q. designed the research, performed experiments, analyzed data, and wrote the manuscript. J.L. performed the FACS analysis, RNA-seq, qPCR and western blot. S. Z. generated AAV vectors. W. Z. and B.W. validated CellREADR AAV vectors in mice. X. A. performed neuron type recording and manipulation experiments in behaving mice. E.A.M, M.A, M.Y and C.P. performed all human CellREADR experiments and analyzed the data. D.G.S. designed and supervised the research involving human tissues, and gave input to the manuscript.

## NOTE

While this manuscript was under revision, the following two preprints were posted on bioRxiv: Kaiyi Jiang, Jeremy Koob, Xi Dawn Chen, Rohan N. Krajeski, Yifan Zhang, Lukas Villiger, Wenyuan Zhou, Omar O. Abudayyeh, Fei Chen, Jonathan S. Gootenberg

Programmable eukaryotic protein expression with RNA sensors https://www.biorxiv.org/content/10.1101/2022.01.26.477951v1

K. Eerik Kaseniit, Noa Katz, Natalie S. Kolber, Connor C. Call, Diego L. Wengier, Will B. Cody, Elizabeth S. Sattely, Xiaojing J. Gao

Modular and programmable RNA sensing using ADAR editing in living cells https://www.biorxiv.org/content/10.1101/2022.01.28.478207v1

## METHODS

### Plasmids

All constructs were generated using standard molecular cloning procedures. Vector backbones were linearized using restriction digestion, and DNA fragment inserts were generated using PCR or gBlock synthesis (IDT). Information on all plasmids is included in **Extended Data Table 1**. All sesRNA inserts are generated by gBlock synthesis (IDT), and sesRNA sequences used in this study are included in **Extended Data Table 2**.

### Cell culture and Transfection

The HeLa cell line was obtained from A. Krainer laboratory (Cold Spring Harbor Laboratory). Mouse neuroblastoma Neuro-2a (N2a) cells were purchased from Millipore-Sigma (Sigma, Cat. 89121404). The HEK293T and KPC1242 cell lines were obtained from D. Fearon laboratory (Cold Spring Harbor Laboratory). HEK293T, Hela and N2a cell lines were cultured in Dulbecco′s Modified Eagle Medium (Corning, 10-013-CV) with 10% fetal bovine serum (FBS) (Gibco, Cat. 16000036) under 5% CO2 at 37□°C. 1% penicillin–streptomycin were supplemented in the medium. Cells were transfected with the Lipofectamine2000 (Invitrogen, Cat. 11668019) DNA transfection reagent according to the manufacturer′s instructions. For interferon treatment, medium containing 1nM Recombinant Mouse IFN-β (R&D, Cat. 8234-MB) was used and changed every 24□hours during the transfection process. For tetracycline treatment, medium containing indicated tetracycline concentration was used 24 hours after transfection to replace the tetracycline free medium and for 48 hours before analysis. For apoptosis assay, cell were incubated for 72 hours after transfection, the RealTime-Glo™ Annexin V Apoptosis and Necrosis Assay (Promega JA1011) was used to quantify the apoptosis level.

### SURVEYOR assay

PCR of Genomic DNA (primers for DYRK1A, Forward: GGAGCTGGTCTGTTGGAGAA; Reverse: TCCCAATCCATAATCCCACGTT) is used to amplify the Cas9 target region from a heterogeneous population of modified and unmodified cells, and the PCR products are reannealed slowly to generate heteroduplexes. The reannealed heteroduplexes are cleaved by SURVEYOR nuclease, whereas homoduplexes are left intact (IDT: Alt-R Genome Editing Detection Kit). Cas9-mediated cleavage efficiency (% indel) is calculated based on the fraction of cleaved DNA^31^.

### Animals

Six to twenty weeks old (both male and female) mice were used in this study. Wide type mice were purchased from Jackson Laboratory (C57BL/6J, 000664). *Fezf2-CreER* and *Rosa26-LoxpSSTOLoxp-H2bGFP* mice were described previously^39,56^. To label *Fezf2^+^* cortical pyramidal neurons, 200mg/kg dose of tamoxifen (T56648, Sigma) was administered by intraperitoneal injection two days before AAV injection. Long Evans rats (*Rattus norvegicus*, aged 12 weeks, 50–250□g) were obtained from Charles River. All the animals were maintained in a 12□h:12□h light:dark cycle with a maximum of five animals per cage for mice and one animal per cage for rats. All animal maintenance and experimental procedures were performed according to the guidelines established by the Cold Spring Harbor Laboratory and Duke Medical Center.

### ADAR1 Knockout cell line construction

ADAR1 Knockout cell line was generated via CRISPR-Cas9 genome editing. The gRNAs were cloned into pSpCas9 (BB)-2A-puro (pX459) (Addgene, Cat. 62988). The gRNA sequences were 5’ GGATACTATTCAAGTCATCTGGG 3’ (gRNA1) and 5’ GTTATTTGAGGCATTTGATG 3’ (gRNA2). Briefly, ADAR1 knockout cells were generated with two gRNAs targeting exon 2, which is shared by both ADAR1-p110 and ADAR1–p150 isoforms. HEK293T cells were seeded into 6 well plates and transfected with 1ug plasmid mixture of pX459-gRNA1 and pX459-gRNA2 using Lipofectamine 2000 (Thermo Fisher Scientific, Cat. 11668019). On the next day, all the media containing transfection reagents was removed and replaced with fresh medium supplemented with puromycin (final conc. 2ug/ml). Fresh medium with puromycin replaced with old medium every two days for three times. The remaining cells after puromycin selection was harvested with TrypLE (Gibco, Cat. 12605036) and distributed in 96-well plate with 1∼2 cells each well. Expanded single clones were screened for ADAR1 deficiency by western blot and disruption of the ADAR1 genomic locus was confirmed by Sanger sequencing.

### Flow cytometry analysis

Two days after transfection, cells were harvested with TrypLE (Gibco, Cat. 12605036), distributed in 96-well round-bottom plate and centrifuged at 500 rpm for 1 min at 4°C. Supernatant was removed and cells were resuspended with 1% PFA buffer and incubated at 4°C overnight or longer time. The cells were then resuspended with 200ul Flow Cytometry Staining Buffer (Invitrogen, Cat. 00-4222-26) before analysis using a LSR Fortessa (BD Biosciences). The Fortessa was operated by FACSDIVA (BD Biosciences) software. Data analysis was performed with FlowJo 10 (FlowJo, LLC). For sorting, cells were submitted to the same procedure as for flow cytometry analysis but without 1% PFA treatment and processed using BD FACSAria (BD Biosciences).

### CellREADR efficiency for exogenous or endogenous transcripts

To assay CellREADR efficiency with fluorescence reporter genes, HEK293T cells or HEK293T ADAR1 knockout cells were first seeded in 24-well plates. After 24□hours, cells were co-transfected with 1.5□μg plasmids in total. At 48 hours of post transfection, cells were collected, fixed with 1% PFA and prepared for FACS analysis. To calculate the CellREADR efficiency from the FACS data, the cells expressing *READR* vector were gated by fluorescent protein (for example, BFP in Fig. 1b-d) expressing by spacer, the efficiency was calculated as the ratio of double positive cell expressing both efRNA and target RNA among all the target RNA expressing cells within the gate. If the spacer is not a fluorescent protein (Fig. 1h and Fig. 2), the ratio was directly calculated as the ratio of double positive cell expressing both efRNA and target RNA among target RNA-expressing cells. For some controls, the CellREADR efficiency was calculated as the percent of efRNA expressing cells among all the counted cells.

### sesRNA design

We used the following procedure for sesRNA design: 1) sesRNA is complementary, i.e. anti-sense, to a specific cellular coding or non-coding RNA sequence; 2) The optimal length of a sesRNA is 200-300 nucleotides; 3) readrRNAconsists of a sesRNA and efRNA arranged in a continuous translation reading frame; 4) One or more STOP codons (TAG) are placed near the center of a sesRNA (ranging between ∼80-220 nucleotide from 5’ end); 5) find 5’-CCA-3’ sequences in the target RNA, which is complementart to a 5’-TGG-3’ sequence in sesRNA; then replace 5’-TGG-3’ with 5’-TAG-3’, so that a mismatch A-C will be introduced when sesRNA base pairs with the target RNA; 6) to ensure that there are no other STOP codons in the sesRNA, all other TAG, TAA, TGA sequences in sesRNA are converted to GAG, GAA, GGA, respectively; preferably the converted STOP codons are not near (i.e. more than10bp from) the TAG defined in step 5; 7) there should be no ATG (initiation codon) after TAG defined in step 5 to exclude the possibility of unintended translation initiation; 8) sesRNAs can be directed to any region of a cellular transcript, including exons, introns, UTR regions, or mature mRNA after splicing. 9) avoid sesRNA with complex secondary structures.

### A-to-I editing rate by CellREADR

To evaluate the A-to-I editing rate, HEK293T cells were seeded in 12-well plates. At 48-hour post transfection, cells were collected and RNA was extracted. RNAs were purified with RNeasy Mini Kit (Qiagen Cat. 74106) and converted to cDNA using TaqMan™ Reverse Transcription Reagents (Thermo Fisher, N8080234). PCR products covering the whole sesRNA region was generated with CloneAmp HiFi PCR Premix (Takara, Cat. No. 639298), and purified with NucleoSpin Gel and PCR Clean-up kit (Takara, Cat. No. 74061) for Sanger sequencing. Editing efficacy was calculated as the ratio of Sanger peak heights G/(A□+□G). Three biological replicates were performed.

### Transcriptome-wide RNA-sequencing analysis

The readrRNA^Ctrl^ or readrRNA^PCNA^, readrRNA^EEF1A1-CDS^-expressing plasmids with the red fluorescent protein (tdTomato) expression cassette were transfected into 293T cells. The RFP^+^ cells (about 5×10^5) were enriched by FACS sorting 48□h after transfection, and RNAs were purified with RNeasy Mini Kit (Qiagen Cat. 74106). To prepare the library, RNA samples were processed with the TrueSeq Stranded mRNA library Prep kit (Illumina, 20020594) and TrueSeq RNA Single Indexes (Illumina, 20020492). Deep sequencing analysis were performed in Illumina NextSeq500 platform at Cold Spring Harbor Laboratory NGS Bioinformatics Center. FastQC was applied to RNA-seq data to check the sequencing quality. All samples passed quality check were mapping to reference genome (GRCh38-hg38). STAR (version 2.7.2) and RSEM (version 1.3.2were used to annotate the genes. TPM values was calculated from RSEM. For all genes with at least one of the libraries above zero reads, the average expression values across biological replicates were compared between samples for detecting differentially expressed genes, using DESeq2^36^ Genes with adjusted p-values < 0.01 and Fold Change > 2 or < 0.5 were identified as significantly differentially expressed. For global A-to-G editing rate analysis, A-to-G editing rate calculation was performed using RNA Editing Detection Pipeline (REDP) which is a python wrapper for REDItool^37^. Parameters were set to default values. Alignment step was done using STAR v2.7.8a with default and coordinate-sorted parameters. Genome reference from Gencode GRCH38.p13 was used and concatenated with known sequences from target genes including *EEF1A1* and *PCNA*.

### Western blot

Primary antibodies against ADAR1 (Santa Cruz, sc-271854, 1:500), GFP antibody (Cell Signaling Technology, Cat. 2956, 1:10000), β-Actin antibody (Cell Signaling Technology, Cat. 3700, 1:10000) were used in this study. We used standard western blot protocols. Briefly, ∼2□×□10^6^ cells were lysed and an equal amount of each lysate was loaded for SDS–PAGE. Then, sample proteins were transferred onto polyvinylidene difluoride membrane (Bio-Rad Laboratories) and immunoblotted with primary antibodies against ADAR1 or GFP, followed by secondary antibody incubation (1:10,000) and exposure.

### Luciferase complementation assay

Forty eight hours after transfection, HEK293T cells expressing firefly luciferase gene were washed in PBS, collected by trituration and transferred to 96-well plates. Promega Luciferase Assay System (Promega, E1500) was used. Firefly luminescence was measured using SpectraMax Multi-Mode Microplate Readers (Molecular Devices).

### qRT-PCR

RNA was extracted and purified with RNeasy Mini Kit (Qiagen Cat. 74106). RNA was converted to cDNA using TaqMan™ Reverse Transcription Reagents (Thermo Fisher, N8080234). qRT-PCR was performed with Taqman probes on the QuantStudio 6 Flex real-time PCR system. The housekeeping gene TBP was used for normalization. The gene probes are purchased from Thermo Fisher Scientific. (Thermo Fisher Scientific, EEF1A1: Hs01591985, PCNA: Hs00427214, XIST: Hs00300535, ACTB: Hs03023943, Mda5: Mm00459183_m1, Rig-I: Mm01216853_m1, Ifnb1: Mm00439552_s1, Il6: Mm00446190_m1, Ccl2: Mm00441242_m1, Fgf2: Mm01285715_m1, Cd40: Mm00441891_m1, Iba1: Mm00479862_g1, Cxcl10: Mm00445235_m1). The housekeeping gene TBP (Thermo Fisher Scientific, human TBP: Hs00427620, mouse Tbp: Mm01277042_m1) was used for normalization.

### Virus production

For producing *READR* and *Reporter* viruses, HEK293T cells were transiently transfected with *READR* or *Reporter* plasmids, AAV serotype plasmids, and pHelper using PEI MAX (catalog 24765, Polysciences Inc.). Seventy-two hours after transfection, the cells were collected in cell culture media, and centrifuged at 4,000rpm for 15 minutes. The supernatant was discarded, the pellet was resuspended in cell lysis buffer, frozen and thawed three times using a dry ice/ethanol bath. Cell lysate was centrifuged at 4,000rpm for 20 minutes. The contaminating DNA in the supernatant was removed by adding Benzonase and was incubated at 37°C for 30 minutes. The viral crude solution was loaded onto an iodixanol density gradient and spun at 60,000 rpm for 90 minutes using a Beckman Ti70 rotor. After the centrifugation, 3-4 ml crude viral preparation was collected from the 40-60% layer with an 18-gauge needle attached to a 10 ml syringe. The viral crude solution was concentrated to 200–250 μl using the Amicon Ultra-15 centrifugal filter (100 kDa), washed with 8 ml PBS once, and concentrated to an appropriate volume. Aliquots were stored at –80°C until use. Mouse and rat *READR* and *Reporter* viruses were packaged as AAV-DJ or AAV-PHP.eB serotypes. *PT enhancer-tTA2* viruses were packaged as PHP.eB with vector from Addgene (#163480). Human READR and Reporter viruses were packaged as AAV2-Retro serotype. *pAAV(DJ)-hSyn-Adar2-sesRNA^Fezf2^-tTA2* and *pAAV(PHP.eB)-hSyn-ADAR2-sesRNA^Ctip2(1)^ -tTA2* were packaged from Vigene Inc.)

### Immunohistochemistry

Mice or rat were anesthetized (using Avertin) and intracardially perfused with saline followed by 4% paraformaldehyde (PFA) in 0.1 M PBS buffer. Following overnight fixation at 4°C, brains were rinsed three times and sectioned 50μm thick with a Leica 1000s vibratome. Sections were placed in blocking solution containing 10% Normal Goat Serum (NGS) and 0.1% Triton-X100 in PBS1X for 1 hour, then incubated with primary antibodies diluted blocking solution overnight at 4 °C. Anti-GFP (1:1000, Aves, GFP-1020); anti-CTIP2 (1:250, Abcam 18465); anti-TLE4 (1:500, Scbt: sc-365406); anti-Rorb (1:300, Novus Biologicals NBP1-82532) were used. Sections were rinsed 3 times in PBS and incubated for 1 h at room temperature with corresponding secondary antibodies (1:500, Life Technologies). Sections were dry-mounted on slides using Fluoromount (Sigma, F4680) mounting medium.

For human tissue, after viral expression had reached a peak, or following patch clamp recording, cortical slices were fixed overnight in 4% PFA, then rinsed in PBS and stored from 1-7 days in PBS with azide. For biocytin recovery, tissue was permeabilized in PBS with 10% TritonX100, 5% NGS, 100uM glycine, and 0.5% BSA for 30 minutes at room temperature on a shaker plate. Both primary and secondary antibodies were incubated 24 hours at 4C. Biocytin was either developed directly with Alexa647 conjugated streptavidin, or indirectly with enzyme metallography method using peroxidase-labeled streptavidin treated with silver ions substrate which deposits metallic silver at the active site. Recovered cells were imaged on a Keyence BZ-X800 with a 10X or 20X objective. Large overview images of native mNeon fluorescence were made on a Leica SP8 upright confocal microscope. For immunohistochemical labeling of FOXP2, and NeuN, tissue was cryoprotected for at least 24 hours in 30% sucrose, then re-sectioned at 40um with a sliding microtome. Floating sections were permeabilized in PBS with 1% TritonX100, 5% NGS, and 100uM glycine for 30 minutes at room temperature and labeled overnight at 4C with primary antibodies (rabbit anti-FOXP2 Abcam #16046 1:500, rat anti-Flag 1:500 Novus NBP1-06712, mouse anti-NeuN Ab 1:500, Cell Signaling #94403S). For secondary antibodies, tissues were either incubated with biotinylated anti-rabbit Ab (1:1000, ThermoFisher 31822) for 2 hours at room temperature followed by streptavidin AlexaFluor647 for 1 hour at room temperature (FOXP2 or with goat anti-mouse Ab (1:1000, ThermoFisher 32723) for 2 hours at room temperature (NeuN).

### In Situ Hybridizaion

All probes were ordered from Molecular Instruments (mouse *Satb2* cat# PRM128, mouse *PlxnD1* cat# PRK885, mouse *Fezf2* cat# PRA339, mouse *vGAT* cat# PRE853, rat *vGAT* cat# PRN024 and human *VGAT* cat# PRM351). Mouse brain was sliced into 50 µm thick slices after PFA perfusion fixation and sucrose protection. Hybridization chain reaction in situ was performed via free floating method in 24 well plate. First, brain slices were exposed to probe hybridization buffer with HCR Probe Set at 37 degree for 24 hours. Brain slices were washed with probe wash buffer, incubated with amplification buffer and amplified at 25 C for 24 hours. On day 3, brain slices were washed, counter stained if needed.

### Stereotaxic viral injection

Adult mice of 8 weeks or older were anesthetized by inhalation of 2% isofluorane delivered with a constant air flow (0.4 L/min). Ketoprofen (5 mg/kg) and dexamethasone (0.5 mg/kg) were administered subcutaneously as preemptive analgesia and to prevent brain edema, respectively, prior to surgery, and lidocaine (2-4 mg/kg) was applied. Mice were mounted in a stereotaxic headframe (Kopf Instruments, 940 series or Leica Biosystems, Angle Two). Stereotactic coordinates were identified. An incision was made over the scalp, a small burr hole drilled in the skull and brain surface exposed. A pulled glass pipette tip of 20–30 μm containing the viral suspension was lowered into the brain; a 500 nl volume of single or mixed viruses was delivered at a rate of 30 nl/min using a Picospritzer (General Valve Corp) into 300 um and 700 um (1:1 volume) of injection sites; the pipette remained in place for 10 min preventing backflow, prior to retraction, after which the incision was closed with nylon suture thread (Ethilon Nylon Suture, Ethicon Inc. Germany) or Tissueglue (3M Vetbond), and animals were kept warm on a heating pad until complete recovery. For rat Tle4 cell type targeting, 3:1 mixture of READR^Tle4^ and Reporter AAVs was stereotactically injected into the cortex (0 mm posterior, 3.5 mm lateral), 1000 ul volume was injected at -2 mm ventral and 1000 ul volume was injected at -1mm ventral, respectively. For rat *vGAT* targeting, 3:1 mixture of *READR^vGAT^* and *Reporter* AAVs was stereotactically injected into the cortex (−4 mm posterior, 3.5 mm lateral), 1000 ul volume was injected at -2.5 mm ventral and 1000 ul volume was injected at -1.5 mm ventral, respectively. Further experiments of immunohistochemistry or in situ hybridization were performed after 3 weeks of virus incubation.

### Physiology in mouse

#### Stereotaxic surgery and viral injection

All surgeries were performed under aseptic conditions and body temperature was maintained with a heating pad. Standard surgical procedures were used for stereotaxic injection and optical fiber implantation. Mice were anesthetized with isoflurane (2-5 % at the beginning and 0.8-1.2 % for the rest of the surgical procedure) and were positioned in a stereotaxic frame and on top of a heating pad maintained at 34-37 °C. Ketoprofen (5 mg/kg) was administered intraperitonially (IP) as analgesia before and after surgery, and lidocaine (2%) was applied subcutaneously under the scalp prior to surgery. Scalp and connective tissue were removed to expose the dorsal surface of the skull. The skin was pushed aside, and the skull surface was cleared using saline. A digital mouse brain atlas was linked to the stereotaxic frame to guide the identification and targeting of different brain areas (Angle Two Stereotaxic System, Leica Biosystems). We used the following coordinates for injections and implantations in the SSp-ul: -0.09 from bregma, 2.46 mm lateral from the midline; CFA: 0.37 mm from bregma, 1.13 mm lateral from the midline.

For viral injection, a small burr hole was drilled in the skull and brain surface was exposed. A pulled glass pipette tip of 20-30 μm containing the viral suspension was lowered into the brain; a 500 nl volume was delivered at a rate of 10-30 nl/min using a Picospritzer (General Valve Corp); the pipette remained in place for 5 min preventing backflow, prior to retraction. Injection was made at a depth of 700 μm. An optical fiber (diameter 200 μm; NA, 0.39) was then implanted in CFA or SSp-ul for optogenetic activation or fiber photometry respectively. The optical fiber was implanted with its tip touching the brain surface. To fix the optical fiber to the skull, a silicone adhesive (Kwik-Sil, WPI) was applied to cover the hole, followed by a layer of dental cement (C&B Metabond, Parkell), then black instant adhesive (Loctite 426), and dental cement (Ortho-Jet, Lang Dental). A titanium head bar was fixed to the skull around lambda using dental cement. Mice were transferred on a heating pad until complete recovery. Further experiments of optogenetic activation or fiber photometry were performed after 8-weeks of virus incubation.

#### In vivo optogenetic activation

We first briefly anesthetized the mice with isoflurane (2%) to attach a reflective marker on the back of their left paws. Mice were then transferred into a tube, head fixed on a stage, and allowed to fully recover from the anesthesia before stimulation starts. A fiber coupled laser (5-ms pulses, 10-20 mW; λ = 473 nm) was used to stimulate at 20 and 50 Hz and constantly for 0.5 s. The inter stimulation interval was 9.5 s. Two cameras (FLIR, FL3-U3-13E4C-C), one frontal and one side camera, were installed on the stage to take high frame rate (100 Hz) videos. The two cameras were synchronized by TTL signals controlled by custom-written MATLAB programs. Videos and TTL-signal states were acquired simultaneously using workflows in Bonsai software. Four LED light lamps were used for illumination (2 for each camera).

#### Behavioral video analysis

The two cameras were calibrated using Camera Calibrator App in MATLAB. For left-paw tracking using MATLAB, the images from the videos were smoothed with a Gaussian lowpass filter (size 9, sigma 1.8). The centroid of the reflective marker on the paw was first detected by a combination of brightness and hue thresholding, then tracked by feature-based tracking algorithms (PointTracker in Computer Vision Toolbox). The tracking results were validated manually, and errors were corrected accordingly. Trials in which mice made spontaneous movements before stimulation onset (within 0.5 s) were excluded from further analysis.

#### Paw stimulation

To stimulate the left paw, mice were lightly anesthetized with isoflurane (0.75-1%). Body temperature was maintained using a feedback-controlled heating pad. A piezo bender (BA4510, PiezoDrive), driven by a miniature piezo driver (PDu100B, PiezoDrive), was insulated with Kapton tape on which a blunt needle was glued. The tip of the needle was attached to the back of the paw to stimulate it with vibration (100 Hz, sine wave, 1 s). Inter stimulation interval was 10 s.

#### In vivo fiber photometry and data analysis

A commercial fiber photometry system (Neurophotometrics) was used to record calcium activities from SSp-ul. A patch cord (fiber core diameter, 200 μm; Doric Lenses) was used to connect the photometry system with the implanted optical fiber. The intensity of the blue light (λ = 470 nm) for GCaMP excitation was adjusted to ∼60 μW at the tip of the patch cord. A violate light (λ = 415 nm, ∼60 μW at the tip) was used to acquire the isosbestic control signal to detect calcium-independent artifacts. Emitted signals were band-pass filtered and focused on the sensor of a CMOS camera. Photometry signal and stimulation onset were aligned based on TTL signals generated by a multifunctional I/O device (PCIe-6321, National Instruments). Mean values of signals from the ROI were calculated and saved by using Bonsai software, and were exported to MATLAB for further analysis.

To process recorded photometry signals, we first performed baseline correction of each signal using the adaptive iteratively reweighted Penalized Least Squares (airPLS) algorithm (https://github.com/zmzhang/airPLS) to remove the slope and low frequency fluctuations in signals. The baseline corrected signals were standardized (Z-score) on a trial-by-trial basis using the median value and standard deviation of 5-s baseline period. The standardized 415-nm excited isosbestic signal was fitted to the standardized 470-nm excited GCaMP signal using robust linear regression. Lastly, the standardized isosbestic signal was scaled using parameters of the linear regression and regressed out from the standardized GCaMP signal to obtain calcium dependent signal.

### Human organotypic sample preparation

Human neocortical tissues (frontal, parietal, temporal) were obtained from pediatirc and adult patients (N=6; ages 6-60) undergoing brain resections for epilepsy. Informed consent was obtained for the donation of tissues under Duke University IRB Protocol 00103019.

Tissue preparation generally followed the methods described in the literature^41,57,58^. Dissection solution contained (in mM): 75 sucrose, 87 NaCl, 25 glucose, 26 NaHCO_3_, 2.5 KCl, 1.2 NaH_2_PO_4_, 10 MgCl_2_, 0.5 CaCl_2_, bubbled with 95/5% O_2_/CO_2_ (pH 7.4). Cortex was carefully dissected under ice cold (∼4C) oxygenated dissection solution to remove the pia. Single gyri were blocked for slicing and cut at 300um thickness on a Leica VT1200. Slices were rinsed in Hank’s Buffered Saline and plated on PTFE membranes (Millipore, cat. # PICMORG50) in 6 well plates. Tissue was cultured in a conditioned media described in the literature^40^. Conditioned media also contained anti-biotic/anti-mycotic (1X, Gibco by ThermoFisher) for the first 7 days *in vitro*, and cyclosporine (5ug/mL) for the entire culture period. HEPES (20mM) was added to the media for the first hour after slicing. AAV retro-grade (AAV-rg) viruses were suspended in conditioned media and applied directly to the slice surface by pipet on the day of plating. Expression of the mNeon reporter was monitored from DIV 3 onward, and tissue was used for immunohistochemistry and patch clamp experiments between DIV 6 and DIV 14.

### Electrophysiology in human organotypic sample

Cultured human neocortical tissues were transferred to a heated recording bath (32-34C) on the platform of an Olympus BX-50 upright microscope. Recording solution contained (in mM): 118 NaCl, 3 KCl, 25 NaHCO_3_, 1 NaH_2_PO_4_, 1 MgCl_2_, 1.5 CaCl_2_, 30 glucose fully saturated with 95/5% O_2_/CO_2_. Borosilicate patch pipettes were pulled with a resistance of 4.0-5.5 MΩ, and filled with an internal solution containing (in mM): 134 K-gluconate, 10 HEPES, 4 ATP-triphosphate Mg salt, 3 GTP-triphosphate Na salt, 14 Phosphocreatine, 6 KCl, 4 NaCl, pH adjusted to 7.4 with KOH. Biocytin (0.2%) was added to the internal solution to allow for morphological identification after recording. Patched cells were held in whole-cell mode for a minimum of 12 minutes to ensure compete filling with biocytin. Cells expressing mNeon were patch-clamped under visual guidance using an Axopatch 700B amplifier (Molecular Devices). Data was digitized with (Digidata 1550B) and recorded with pClamp 10 (Molecular Devices). Membrane voltage was recorded at 100k Hz and low-pass filtered at 10k Hz. Liquid junction potential was not corrected. Pipette capacitance and series resistance were compensated at the start of each recording, and checked periodically for stability of the recording configuration. Intrinsic membrane properties were measured with a -10pA current step. Firing rheobase was measured with a ramp current of 1s duration and 100-300pA final amplitude. Input-output curves were generated from a series of current step starting at -50pA and increasing in 10pA increments until a maximum firing rate was elicited. Data were analyzed using NeuroMatic and in-house routines in Igor Pro (Wavemetrics).

### Microscopy and Image Analysis

Cell imaging in tissue culture was performed on ZEISS Axio Observer (CSHL St. Giles Advanced Microscopy Center). Imaging from serially mounted brain sections was performed on a Zeiss LSM 780 or 710 confocal microscope (CSHL St. Giles Advanced Microscopy Center) and Nikon Eclipse 90i fluorescence microscope, using objectives X63 and x5 for embryonic tissue, and x20 for adult tissue, as well as x5 on a Zeiss Axioimager M2 System equipped with MBF Neurolucida Software (MBF). Quantification and image analysis was performed using Image J/FIJI software. Statistics and plotting of graphs were done using GraphPad Prism 9.

### Statistics

Unpaired two-tailed Student’s t-test was used for group comparison. Statistical analyses were performed Prism 9 (GraphPad Software, Inc.). DESeq2 was used to analyze statistical significance of transcriptome-wide RNA-seq data.

---

**Extended Data Fig. 1.**
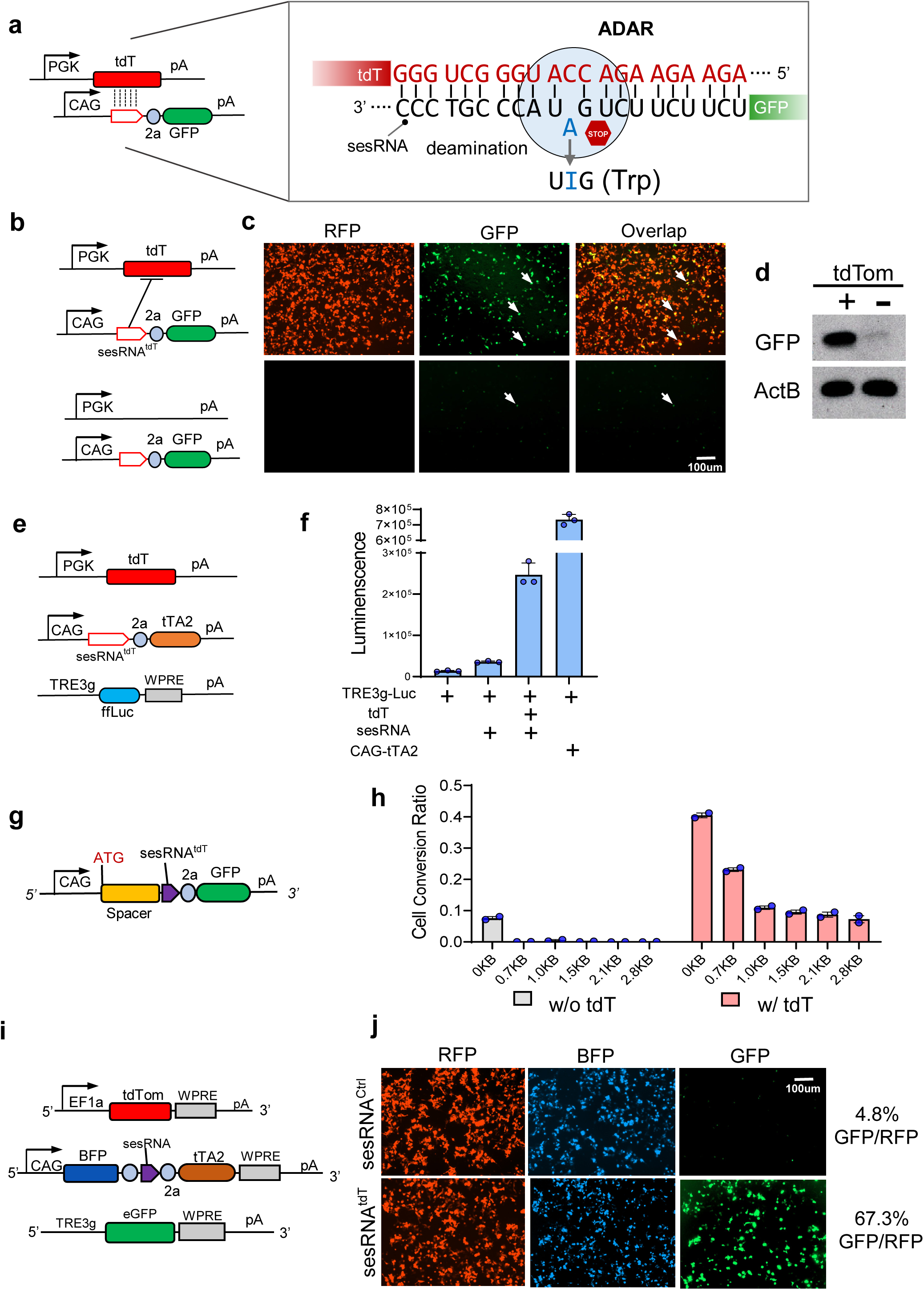
Design and test of singular and binary CellREADR vectors. **a,** Schematic of a singular CellREADR vector. Left, *PGK*-*tdT* expresses the tdTomato target RNA from a PGK promoter. *READR^tdT-GFP^*expresses a READR RNA consisting of sesRNA^tdT^ and efRNA^GFP^, driven by a CAG promotor. Vertical dashed lines indicate the complementary base pairing region between tdT mRNA and sesRNA^tdT^, with sequence surrounding the editable STOP codon shown on the right. At the editing site, the editable adenine in sesRNA^tdT^ (cyan) is mismatched to a cytosine in the tdT mRNA. TdTomato is a tandem repeat of two dTomato genes, thus a tdT RNA contains two copies of target sequence for sesRNA^tdT^ base pairing. **b-d,** Validation of the *READR^tdT-GFP^* vector. In 293T cells co-transfected with *READR^tdT-GFP^*and *PGK*-*tdT* **(b)**, many switched on GFP translation and fluorescence (**c**, upper arrows). In cells co-transfected with control empty vector, few cells showed GFP expression (**c**, lower). GFP expression was further assayed by Western blotting (**d**). **e,** A binary vector design for CellREADR luciferase assay. *READR^tdT-tTA2^*expresses a readrRNA consisting of sesRNA^tdT^ and efRNA^tTA2^, and *TRE-ffLuc* expresses the luciferase RNA upon tTA2 activation. **f,** Luciferase activity dramatically increased only in cells transfected with three vectors in (**e**). Co-transfection of *TRE-ffLuc* with *CAG-tTA2*, which constitutively expresses tTA2, served as a positive control. **g,** Schematic *READR^tdT-GFP^* vector in which a spacer sequences is inserted before *sesRNA^tdT^* coding region. **h,** 293T cells were transfected with *READR^tdT-GFP^* vector encoding viable length of spacers without (gray) or with (pink) tdT target RNA expression, respectively. Quantification of conversion ratio calculated as percentage of GFP^+^ cells among RFP^+^ cells. **i,** A binary vector design for CellREADR assay. **j,** Representative images of GFP conversion with binary vectors in (**i**). In cells co-transfected with sesRNA^Ctrl^ vector, few GFP^+^ cells were observed. Conversion percentages are shown on the right. Error bars in **f** and **h** are mean values□±□s.e.m. n□=□3 for **f**, n = 2 for **h**, n represents the number of independent experiments performed in parallel.

**Extended Data Fig. 2.**
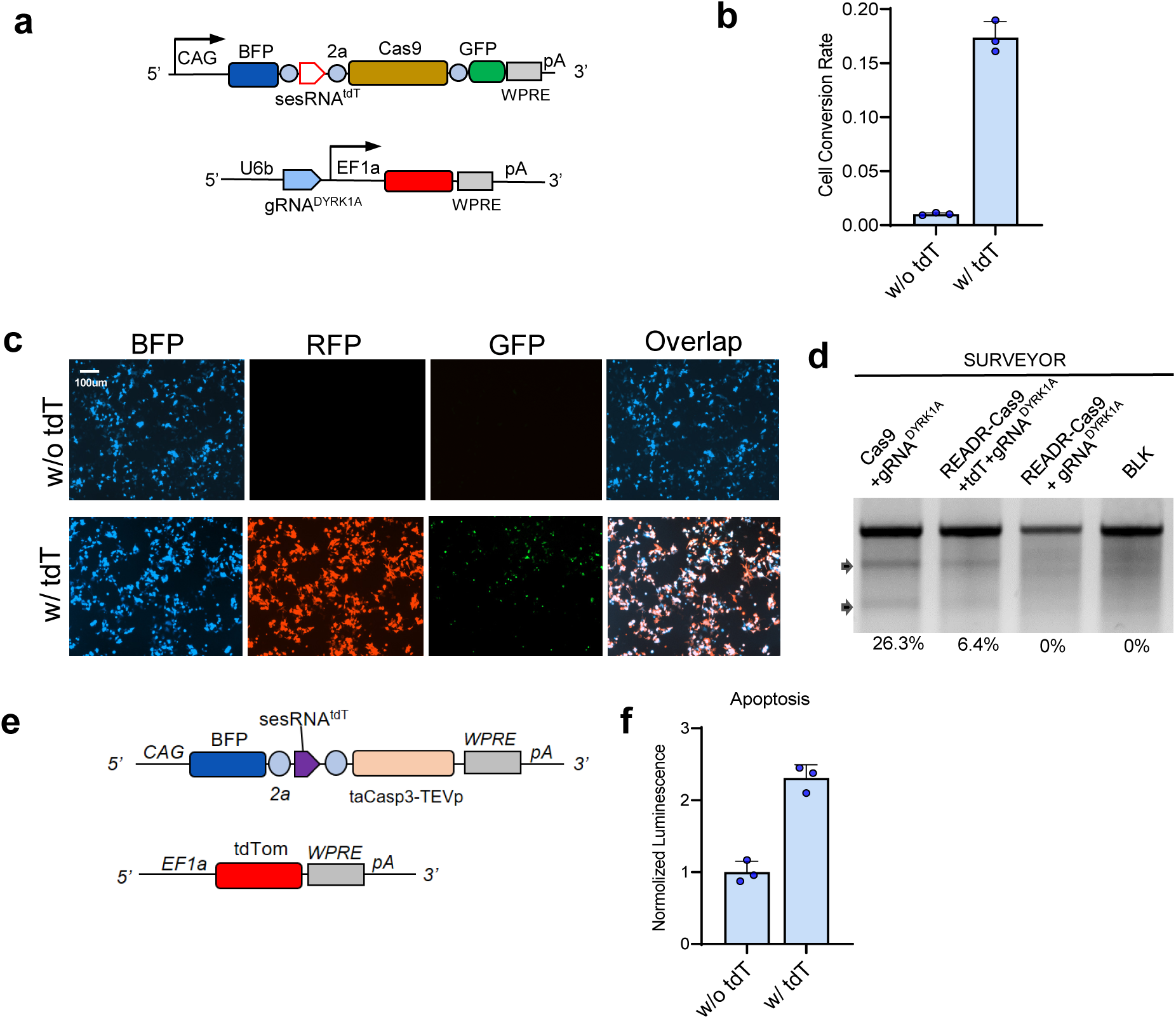
CellREADR enables RNA sensing dependent gene editing and cell ablation. **a,** Vector design for CellREADR-mediated and target RNA-dependent gene editing. In *READR ^tdT-Cas9/GFP^*, a CAG promoter drives expression of BFP followed by sequences coding for *sesRNA^tdT^*, Cas9, and eGFP effectors. In another vector, EF1a promoter drives tdT expression and U6b promoter drives the expression of a guide RNA (gRNA) targeting the DYRK1A gene in 293T cells. **b,** Quantification of *READR^tdT-Cas9/GFP^* efficiency as percent of GFP among RFP and BFP expressing cells with or without tdT target RNA. **c,** Cells transfected with the both *U6b-gRNA^DYRK1A^-CAG-tdT* and *READR^tdT-Cas9/GFP^*showed robust GFP expression co-localized with BFP and RFP (bottom). Cells transfected with *READR ^tdT-Cas9/GFP^* only (top) showed almost no GFP expression. **d,** SURVEYOR assay showed Cas9-mediated cleavage in the human DYRK1A locus. DNA cleavage was observed in cell lysates transfected with *U6b-gRNA^DYRK1A^-CAG-tdT* and *READR ^tdT-Cas9/GFP^*, but not in *U6b-gRNA^DYRK1A^* and *READR^tdT-Cas9/GFP^* that lacked tdT target RNA. CAG-Cas9 with *U6b-gRNA^DYRK1A^* cell lysate and 293T cell lysate without plasmid transfection were used as positive control and negative control, respectively. Arrows indicate cleavage products. **e,** Vector design for CellREADR-mediated and target RNA-dependent cell death induction. In *READR ^tdT-taCasp3-TEVp^*, a CAG promoter drives expression of BFP followed by sequences coding for sesRNA^tdT^ and taCasp3-TEVp as effector to induce cell death. **f.** Cell apoptosis level measured by luminescence was increased in the cells transfected *READR ^tdT-taCasp3-TEVp^* and *EF1a-tdT* compared with cells with no tdT RNA. Error bars in **b** and **f** are mean values□±□s.e.m. n□=□3, n represents the number of independent experiments performed in parallel.

**Extended Data Fig. 3.**
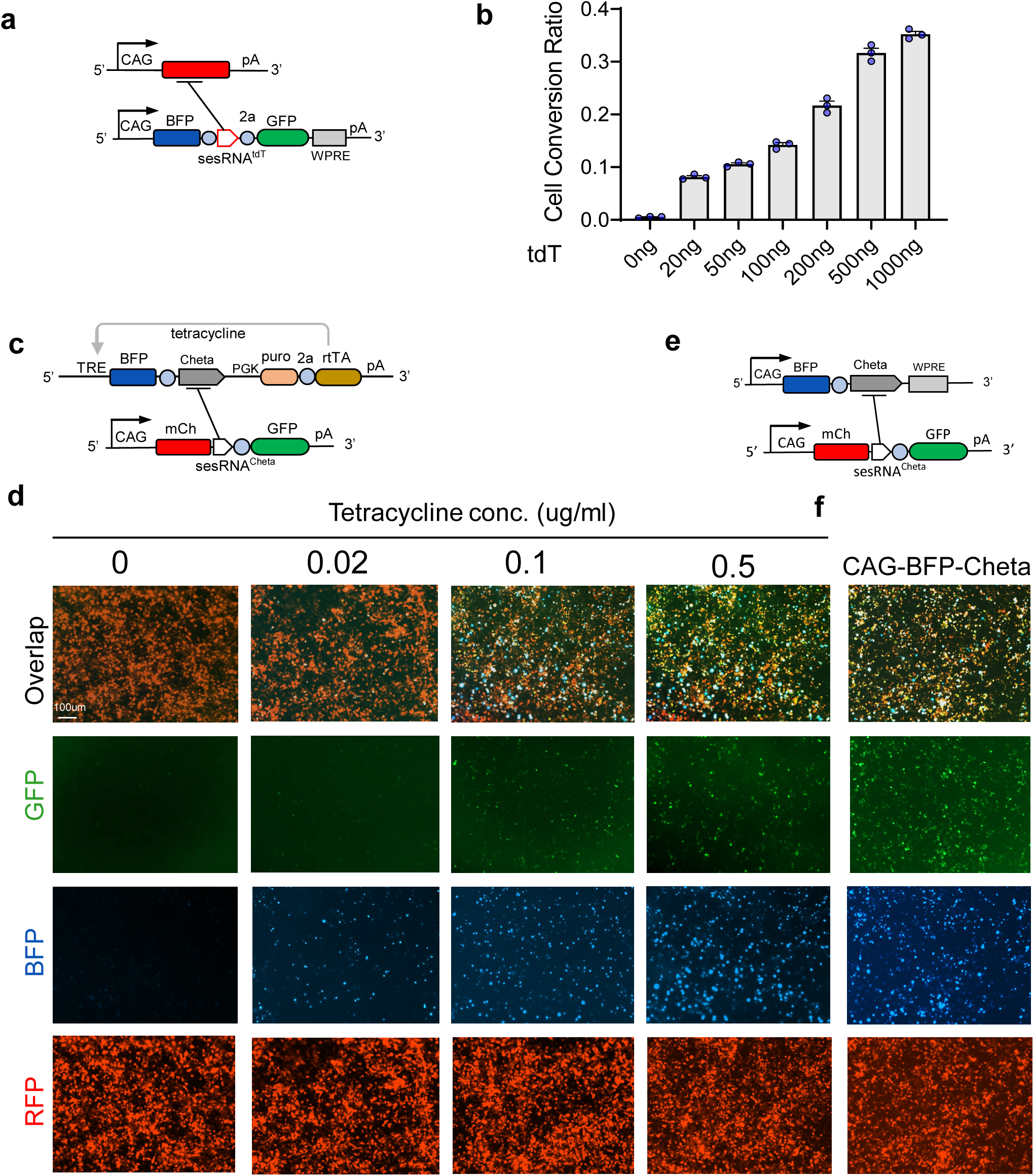
CellREADR efficiency correlates with the expression level of target RNA. **a,** Vector design of a tri-color CellREADR assay system. **b,** Quantification of *READR^tdT-GFP^* efficiency with increasing amount of *CAG-tdT* vector used for co-transfection of 293T cells. **c,** Schematics of *rtTA-TRE-ChETA* and *READR^ChETA-GFP^* vectors (also see **Fig. 1e**). **d,** Representative images of co-transfected 293Tcells treated with different concentration of tetracycline in culture medium (also see **Fig. 1e, f, g**). **e-f,** Co-transfection of *READR^ChETA-GFP^* with a vector that constitutively expresses ChETA (**e**) resulted highest conversion to GFP expression cells (**f**) compared to those in (**d**). Error bars in **b** are mean values□±□s.e.m. n□=□3, n represents the number of independent experiments performed in parallel.

**Extended Data Fig. 4.**
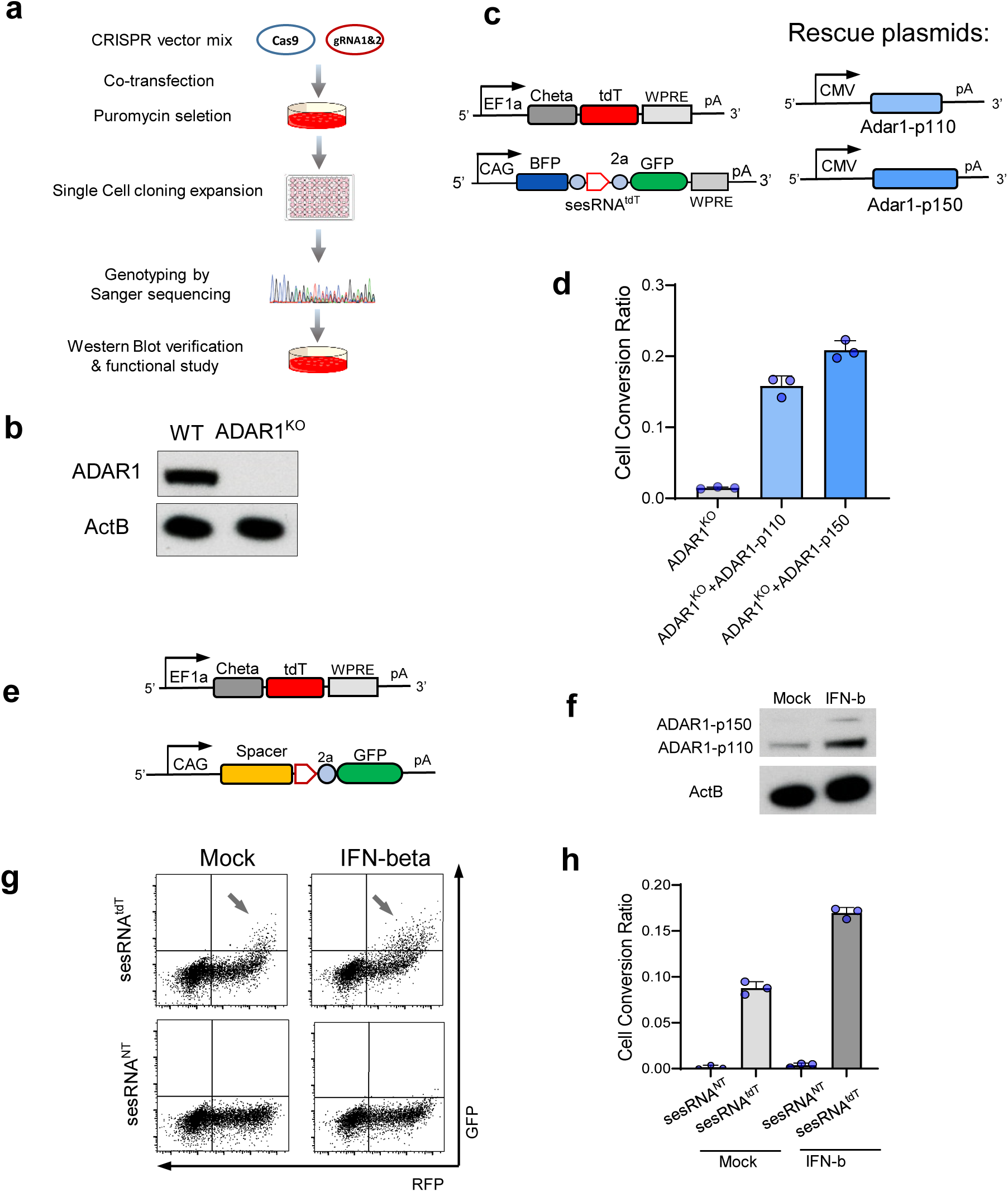
ADAR1 is necessary for CellREADR in 293T cells. **a,** Schematic for generating a ADAR1 knockout cell line with CRISPR/Cas9. **b,** Western blot analysis showing ADAR1 expression in wild-type and no expression in ADAR1 knockout cells. **c,** Schematics of *EF1a-ChETA-tdT* and *READR^tdT-GFP^* vectors (left), and ADAR1 isoform expression vectors (right). **d,** Both p110 and p150 ADAR1 isoforms rescued the CellREADR functionality assayed by cell conversion ratio in ADAR1 knockout cells. **e-h**, INF-b increased CellREADR efficiency and ADAR expression. **e,** Schematic of *EF1a-ChETA-tdT* and *READR^tdT-GFP^* vectors. **f,** Western blot analysis showed increased expression of ADAR1-p110 protein and induction of ADAR1-p150 isoform after interferon treatment. **g,** Representative FACS analysis of GFP and RFP expression with mock (left) or interferon treatment (right). **h,** Quantification of the *READR^tdT-GFP^*efficiency in (**g**), which was increased by interferon treatment. Error bars in **f** and **d** and **h** are mean values□±□s.e.m. n□=□3, n represents the number of independent experiments performed in parallel.

**Extended Data Fig. 5.**
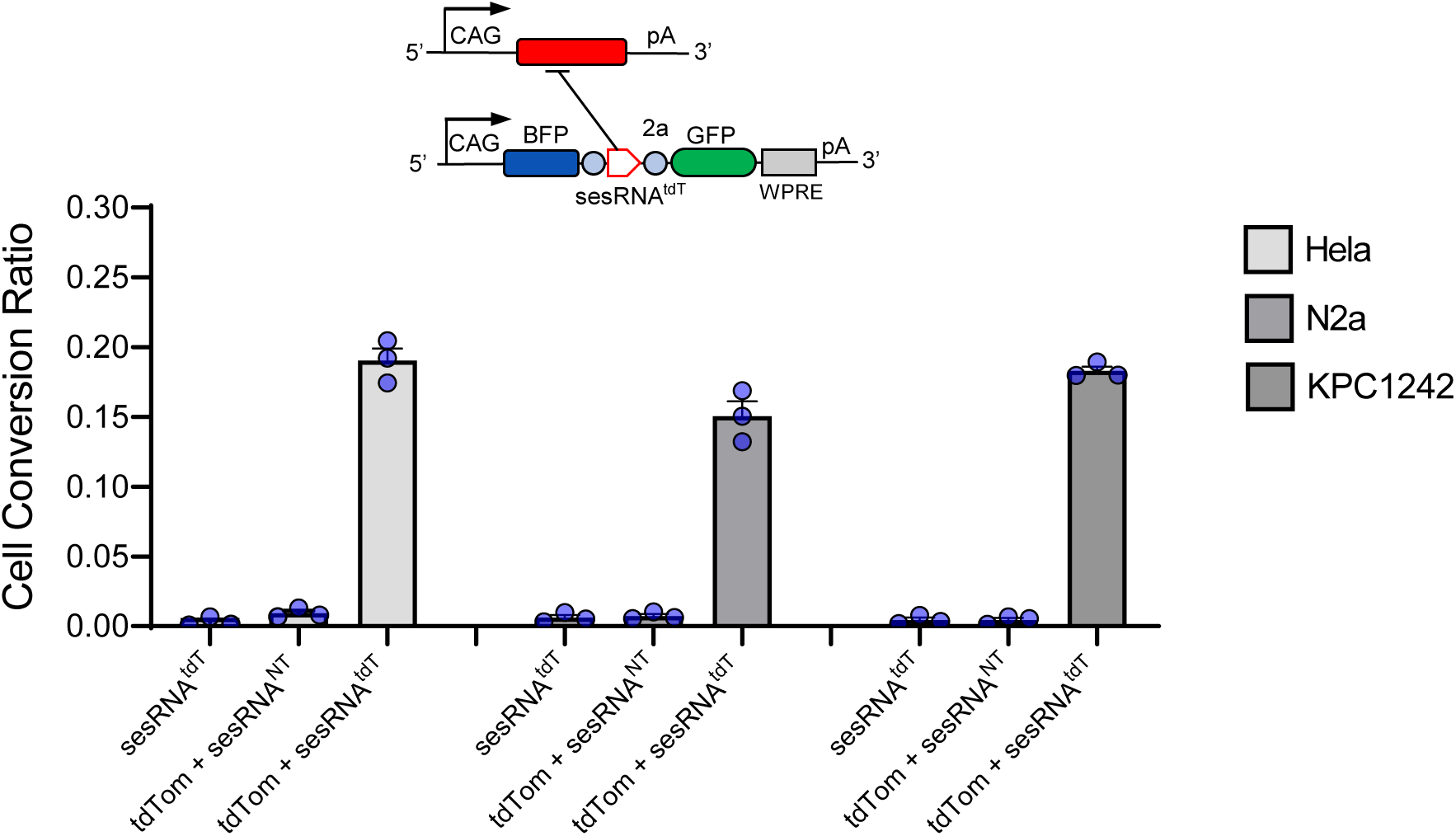
CellREADR functions in multiple human and mouse cell lines. Schematic of CellREADR vectors. *CAG-tdT* expresses the tdTomato target RNA from a CAG promoter. *READR^tdT-GFP^* expresses a READR RNA consisting of sesRNA^tdT^ and efRNA^GFP^, driven by a CAG promotor (top). Hela, N2a, KPC1242 cell lines with comparable efficiency from human (Hela) and mice (N2a, KPC1242) (bottom). Error bars are mean values□±□s.e.m. n□=□3, n represents the number of independent experiments performed in parallel.

**Extended Data Fig. 6.**
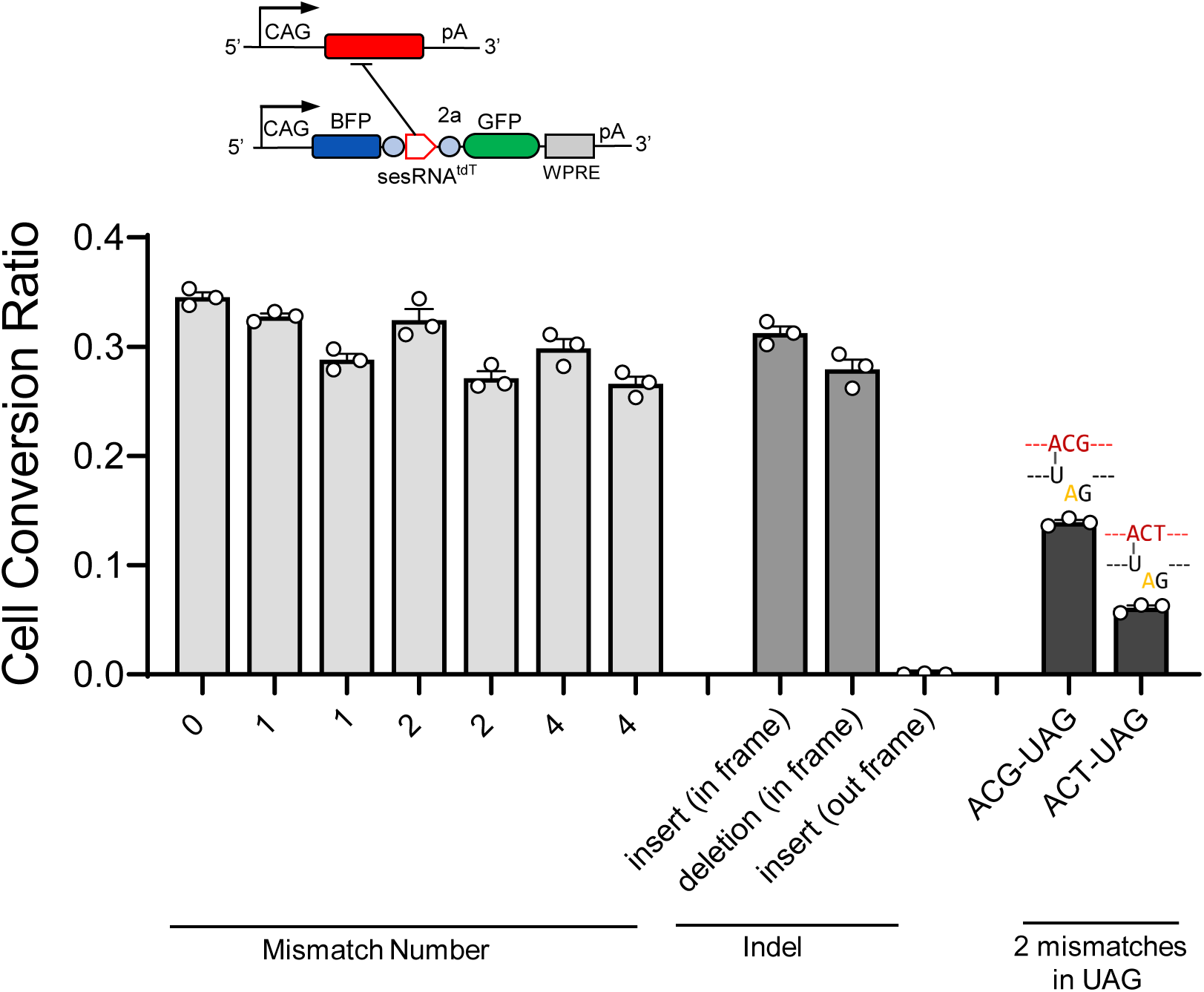
Effects of different nucleotide mismatches between sesRNA and target RNA. Schematic of CellREADR vectors (top, related to **Figure 2b**). CellREADR efficiency measured as RFP to GFP conversion ratio with different types of mismatch number, indel or two mismatches within UAG STOP codon. Error bars are mean values□±□s.e.m. n□=□3, n represents the number of independent experiments performed in parallel.

**Extended Data Fig. 7.**
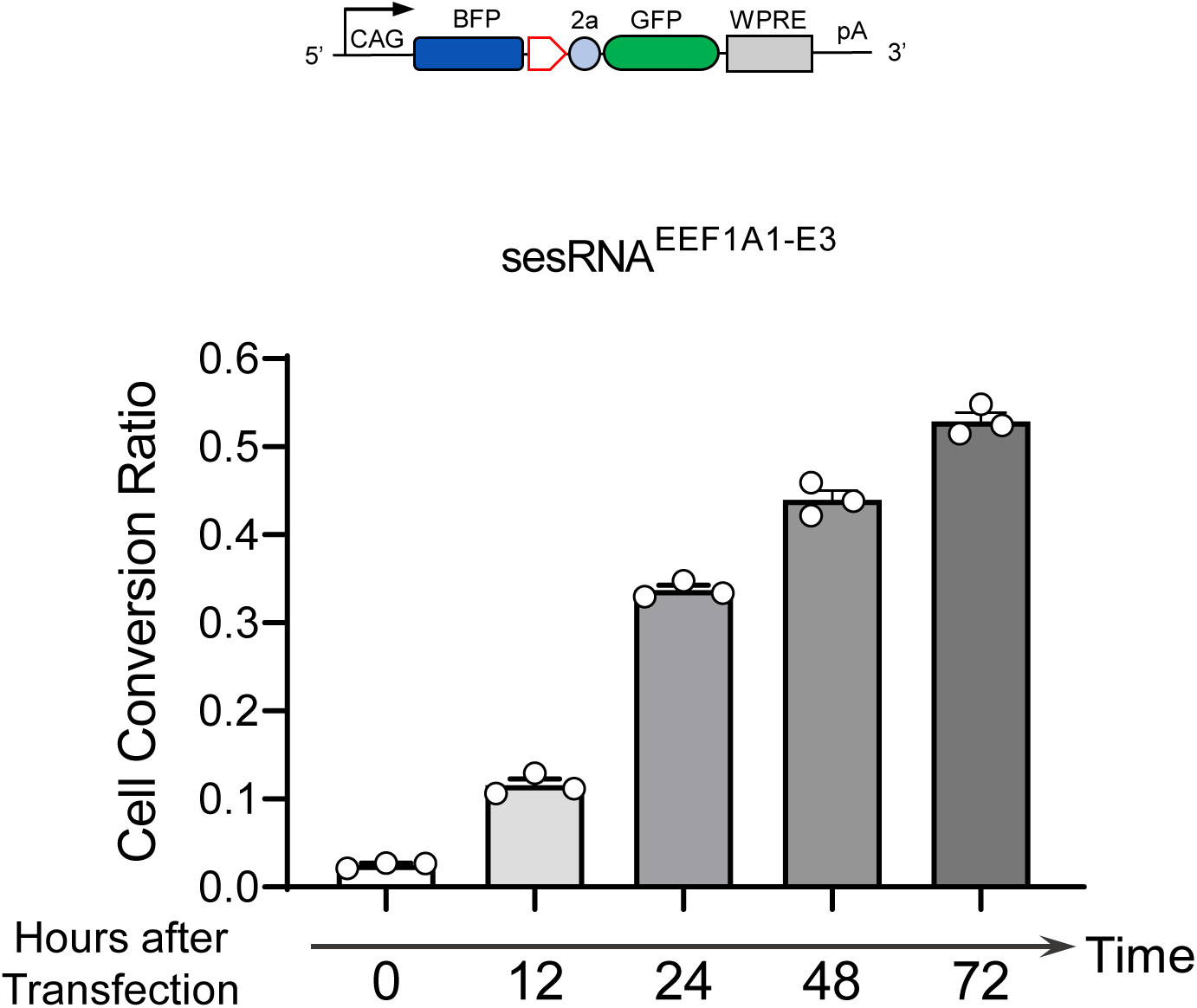
CellREADR mediated RNA sensing and effector translation accumulate with time. CellREADR mediated sensing of endogenous EEF1A1 mRNA and effector translation in 293T cells increased with longer incubation time after transfection. Error bars are mean values□±□s.e.m. n□=□3, n represents the number of independent experiments performed in parallel.

**Extended Data Fig. 8.**
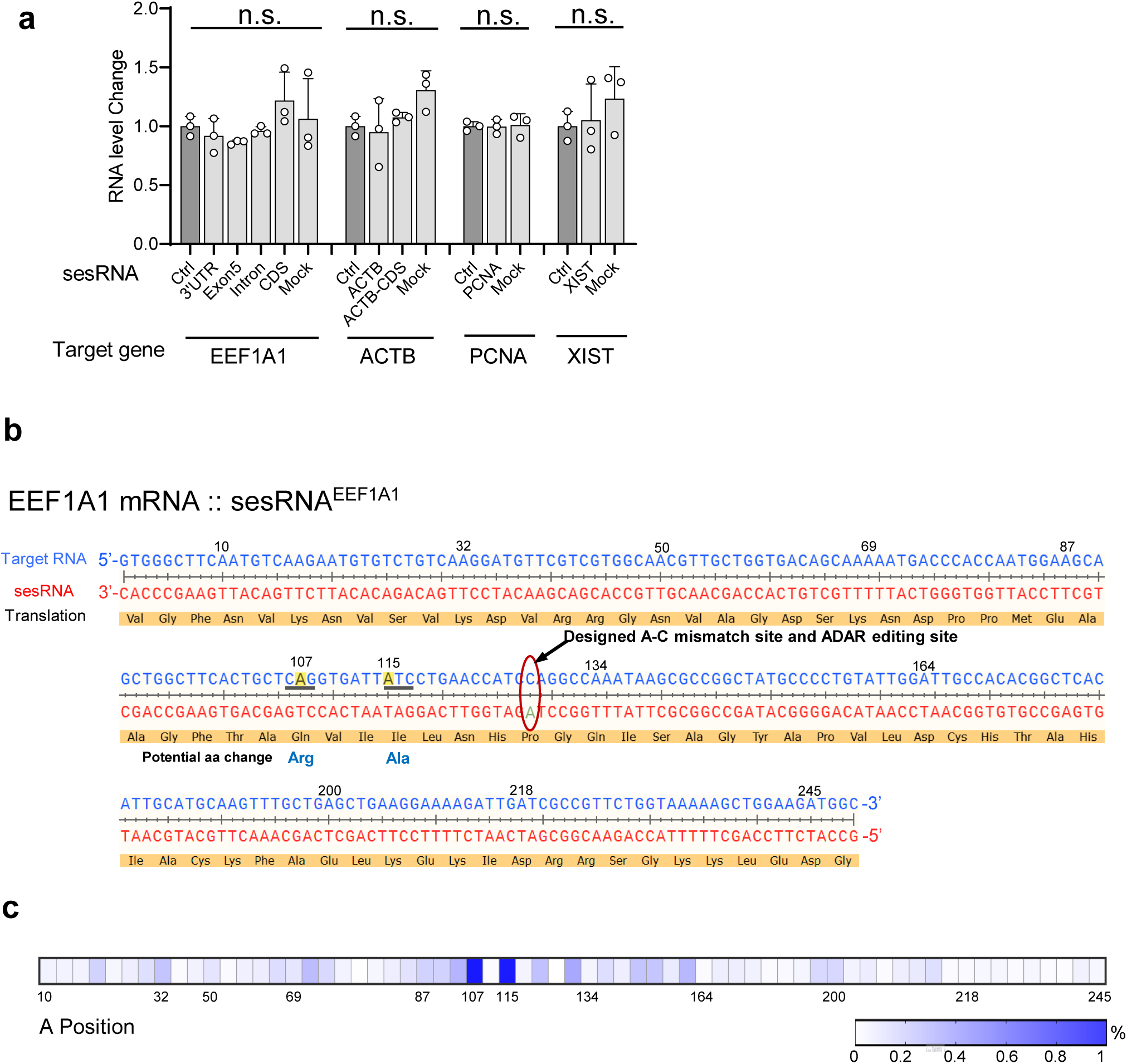
Effects of CellREADR on targeted mRNA. **a,** Quantitative PCR showing that CellREADR-mediated sesRNA expression did not impact the expression levels of targeted RNAs. **b,** Base pairing of EEF1A1 mRNA and sesRNA^EEF1A1-CDS^. EEF1A1 mRNA or sesRNA was represented in blue and red, respectively. Peptide translated from EEF1A1 mRNA was highlighted in brown. Targeted region of EEF1A1 mRNA was analyzed by RNAseq. **c**, The ratios of A-to-G changes in EEF1A1 mRNA at each adenosine position was quantified and shown in heatmap. Two adenosines (A107 and A115) showed higher rate of A-to-G editing (**c**). Off-target editing of two sensitive adenosines can induce potential animo acid change (underlined in **b**). Error bars are mean values□±□s.e.m. n = 3, n represents the number of biological replicates performed.

**Extended Data Fig. 9.**
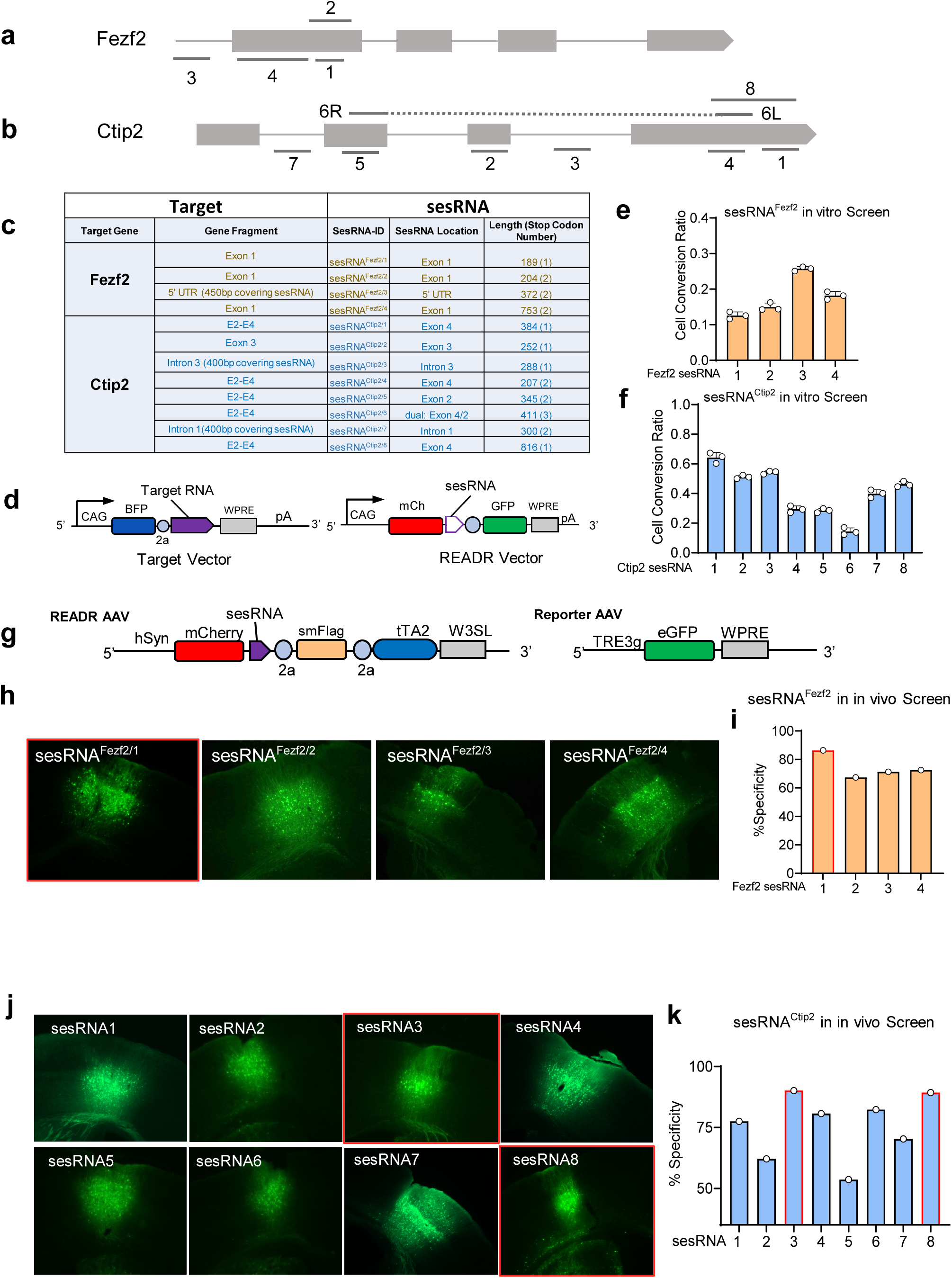
Design and screen of sesRNAs targeting Fezf2 and Ctip2 RNAs in vitro and in vivo. **a-b,** Genomic structures of mouse Fezf2 (**a**) and Ctip2 (**b**) genes with locations of various sesRNAs as indicated. **c,** List of sesRNAs and Fezf2 and Ctip2 target gene fragments used for sesRNA screen. **d,** In Target vectors *CAG-BFP-Fezf2* or *CAG-BFP-Ctip2*, a 200-3000 bp genomic region of the Fezf2 or Ctip2 gene containing sequences complementary to a sesRNA in (**a, b**) were cloned downstream to the BFP and T2a coding region driven by a CAG promoter. In *READR* vectors, *READR^Fezf2-GFP^* or *READR^Ctip2-GFP^*expresses corresponding sesRNAs shown in (**a, b**). **e-f,** Quantification of efficiencies *READR^Fezf2-GFP^* (**e**) or *READR^Ctip2-GFP^*(**f**) as GFP conversion ratio by FACS assay of 293T cells co-transfected with *CAG-BFP-Fezf2* or *CAG-BFP-Ctip2* target vector, respectively. **g,** Schematic of binary *READR* AAV vectors. In *READR* vector, a hSyn promoter drives expression of mCherry followed by sequences coding for sesRNA^Ctip2^, smFlag and tTA2 effectors. In Reporter vector, TRE promoter drives mNeonGreen in response to tTA2 from the *READR* vector. **h,** Coronal sections of mouse cortex injected with binary *READR^Fezf2^* vectors. mNeonG indicated *READR^Fezf2^* labeled cells. Four *Fezf2* sesRNAs were screened. **i,** Quantification of specificity of 4 *Fezf2* sesRNAs in (**h**). For Fezf2 sesRNA in-vivo screen, the specificity of each sesRNA was calculated by co-labeling by *READR* AAVs and CTIP2 antibody (due to lack of FEZF2 antibody); as Ctip2 represents a subset of Fezf2+ cells (not shown), CTIP2 antibody gives an underestimate of the specificity of Fezf2 sesRNA. SesRNA1 showed highest specificity. **j,** Coronal sections of mouse cortex injected with binary *READR^Ctip2^* vectors. mNeonG indicated binary *READR* labeled cells. Eight *Ctip2* sesRNAs were screened. **k,** Quantification of specificity of 8 sesRNAs in (**j**). The specificity of each sesRNA was calculated by co-labeling by binary *READR^Ctip2^*AAVs and CTIP2 antibody (not shown). SesRNA3 and sesRNA8 showed highest specificity.

**Extended Data Fig 10.**
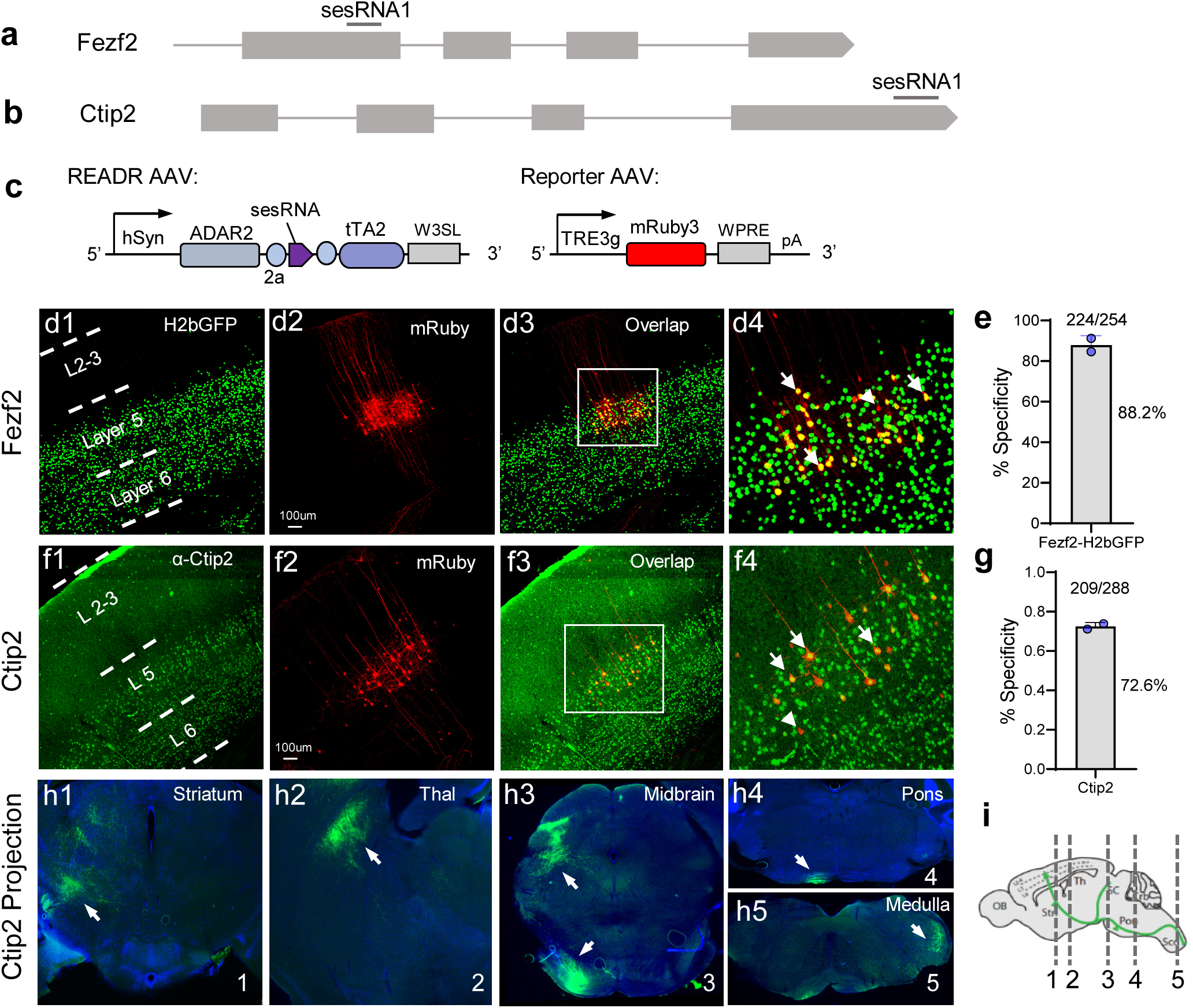
CellREADR targeting of PN^Fezf2^ and PN^Ctip2^ types in mouse cortex with ADAR2 overexpression. **a-b,** Genomic structures of mouse *Fezf2* (**a**) and *Ctip2* (**b**) genes with locations of sesRNAs as indicated, respectively. **c,** Schematic of binary AAV vectors for targeting neuron types. In *READR^Fezf2-tTA2^*, a hSyn promoter drives expression of ADAR2 followed by sequences coding for sesRNA^Fezf2(1)^, T2a, and tTA2 effector. In *TRE-mRuby*, the TRE promoter drives mRuby3 in response to tTA2 from the *READR* virus. **d,** Image of coronal section from a *Fezf2-CreER;LoxpSTOPLoxp-H2bGFP* mouse brain, showing the distribution pattern of *Fezf*2^+^ PNs in S1 somatosensory cortex (**d1**). Co-injection of AAVs *READR^Fezf2^*^(1)*-tTA2*^ and *TRE-mRuby* specifically labeled PNs in L5b and L6 (**d2)**. Co-labeling by CellREADR AAVs and H2bGFP (**d3**) with magnified view in (**d4**). Arrows indicate co-labeled cells; arrowhead shows a neuron labeled by CellREADR AAVs but not by *Fezf2-H2bGFP* (**d4**). **e,** Specificity of *READR^Fezf2(1)-tTA2^*. **f,** Coronal section of WT brain immuno-stained with a CTIP2 antibody, showing the distribution pattern of Ctip2^+^ PNs in S1 cortex (**f1**). Co-injection of AAVs *READR^Ctip2(1)^* and *TRE-mRuby* specifically labeled PNs in L5b (**f2**). Co-labeling by CellREADR AAVs and CTIP2 antibody (**f3**) with magnified view in (**f4**). Arrows show the co-labeled cells; arrowhead showed a mis-labeled cells by *READR^Ctip2(1)^*(**f4**). **g,** Specificity of *READR^Ctip2(1)^*. **h,** Axonal projection pattern of AAV *READR^Ctip2(1)-tTA2^* and *TRE-eYFP* infected PNs in S1 cortex. Representative images showing projections to striatum, thalamus, midbrain, pons and medulla (arrows). **i,** Schematic locations of coronal sections are shown on the right panel.

**Extended Data Fig. 11.**
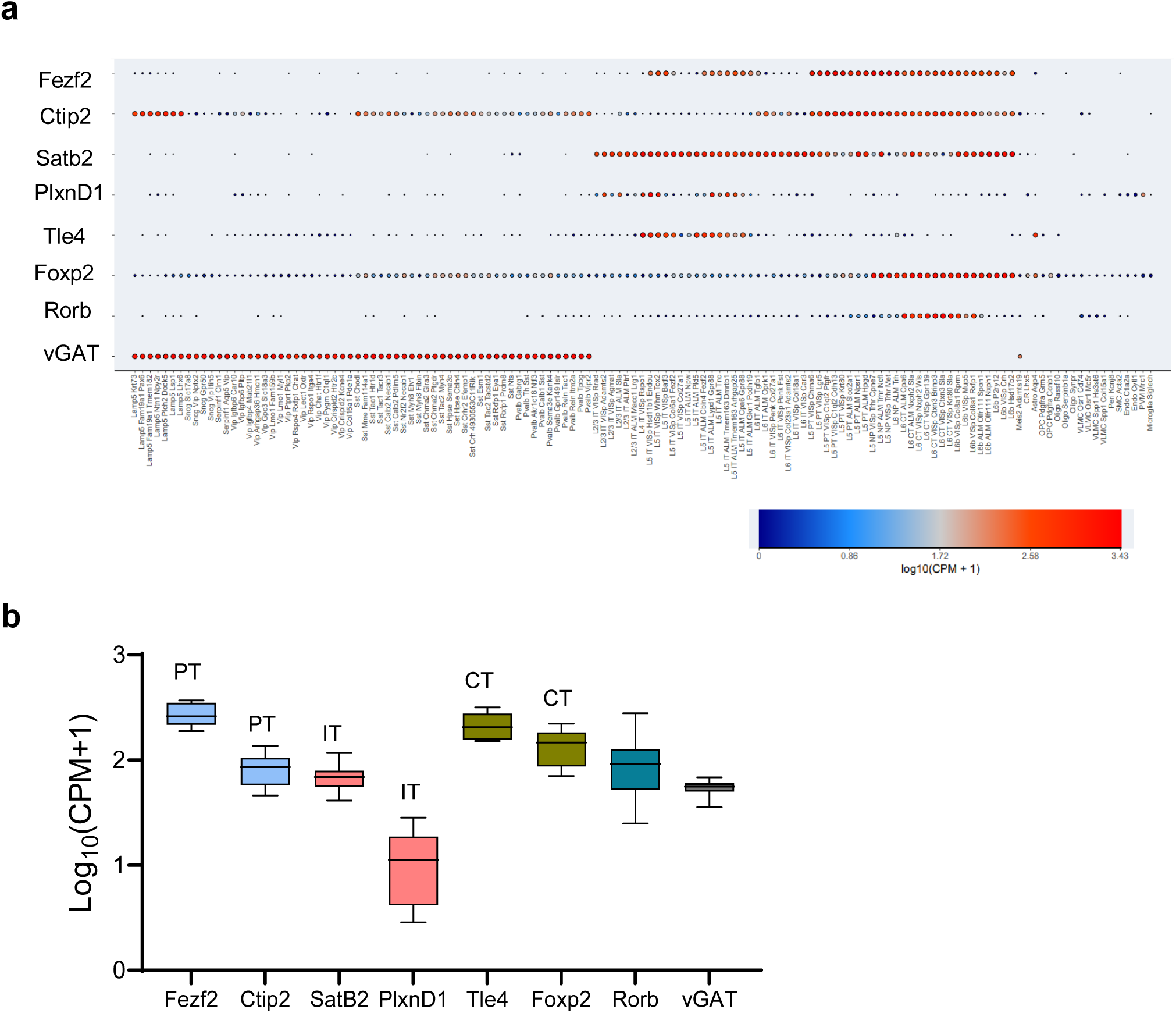
Expression level and laminar distribution of cortical cell type markers in mice. **a.** Group plot of selected genes in transcriptomic cell type clusters, based on dataset from the Allen Institute for Brain Science. Gene expression level and cortical distribution were shown. **b,** gene expression level of major cell type marker genes. The plots were generated with scRNAseq data^40^.

**Extended Data Fig. 12.**
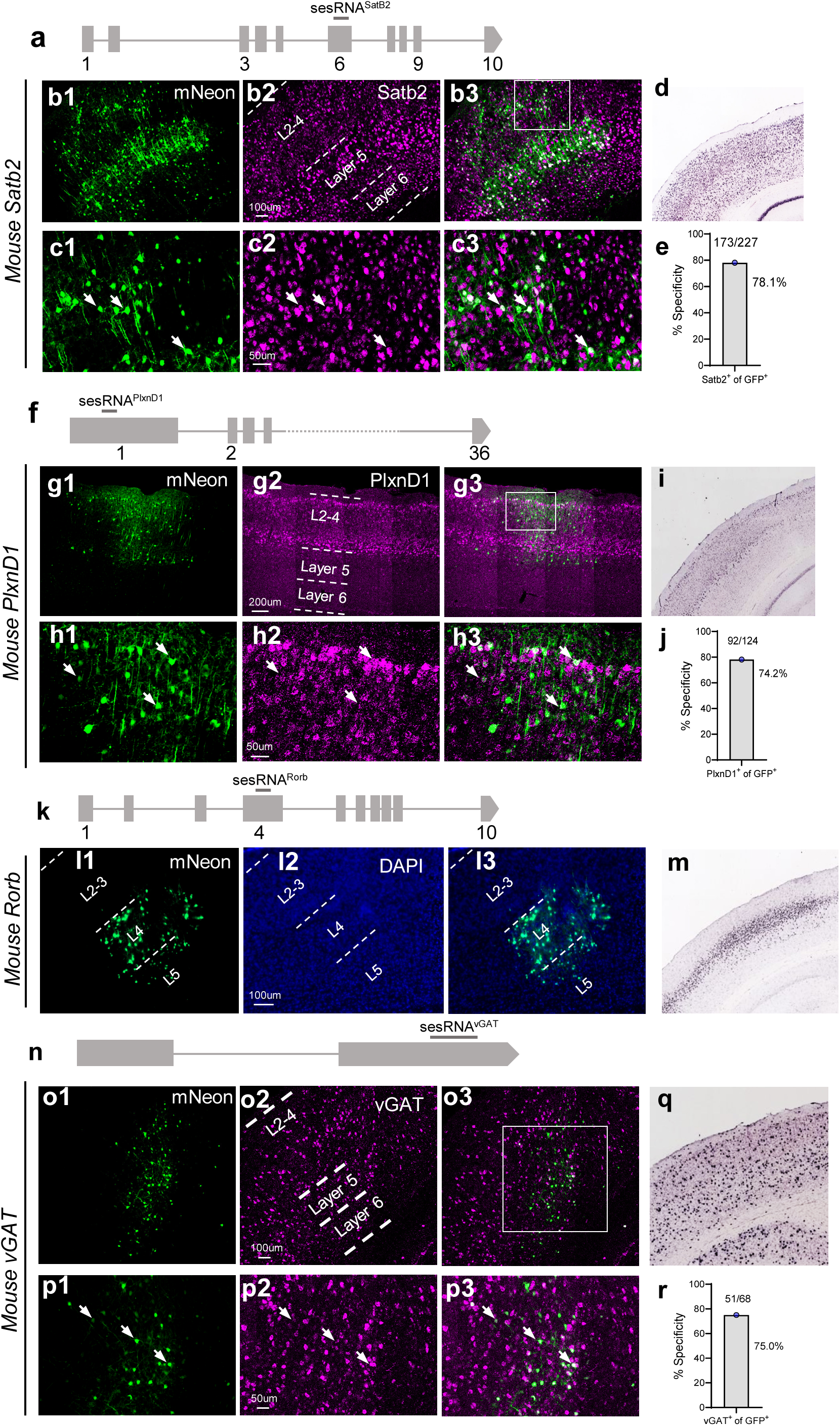
CellREADR targeting of additional cortical neuron types in the mouse. **a,** Genomic structures of mouse the *Satb2* gene with location of a sesRNA as indicated. **b-c,** Cell labeling pattern in S1 by co-injection of binary vectors described in **Fig 4i**. (**b1**) AAVs *READR^Satb2^*and *TRE-mNeon* labeled cells in both upper and deep layers. (**b2**) *Satb2* mRNA in-situ hybridization. (**b3**) Co-labeling by *READR^Satb2^* and *Satb2* mRNA. **c,** Magnified view of boxed region in (**b3**). Arrows indicate co-labeled cells. **d,** *Satb2* mRNA expression pattern in S1 cortex at P56 from the Allen Mouse Brain Atlas. **e,** Specificity of *READR^Satb2^*measured as the percent of *Satb2^+^* cells among mNeon cells. **f,** Genomic structures of the mouse *PlxnD1* gene with location of a sesRNA as indicated. **g-h,** AAVs *READR^PlxnD1^* and *TRE-mNeon* labeled cells in upper layers and L5a in S1 (**g1**). *PlxnD1* mRNA in-situ hybridization (**g2**). Co-labeling by *READR^PlxnD1^* AAVs and *PlxnD1* mRNA (**g3**). **h,** Magnified view of boxed region in (**g3**). Arrows show the co-labeled cells. **i,** *PlxnD1* mRNA expression in P56 S1 cortex from the Allen Mouse Brain Atlas. **j,** Specificity of *READR^PlxnD1^* measured as the percent of *PlxnD1^+^* cells among mNeon cells. **k,** Genomic structures of the mouse *Rorb* gene with location of a sesRNA as indicated. **l,** AAVs *READR^Rorb^*and *TRE-mNeon* labeled cells in layer 4 (**l1, l3**). DAPI staining indicated laminar structure (**l2**). mNeon labeling pattern is consistent with *Rorb* mRNA expression in P56 S1 cortex from the Allen Mouse Brain Atlas (m); the specificity of Rorb sesRNA is yet to be rigorously quantified due to the lack of a specific antibody to RORB and a in situ probe to *Rorb* mRNA. **n,** Genomic structures of mouse the *vGAT* gene with location of a sesRNA as indicated. **o-p,** binary *READR^vGAT^* and *TRE-mNeon* labeled cells (**o1**). *vGAT* mRNA in-situ hybridization. (**o2**). Co-labeling by *READR^vGAT^* AAVs and *vGAT* mRNA (**o3**). **p,** Magnified view of rectangle in (**o3**). Arrows show t, he co-labeled cells. **q,** *vGAT* mRNA expression in P56 S1 cortex from the Allen Mouse Brain Atlas. **r,** Specificity of *READR^vGAT^* measured as the percent of *vGAT^+^* cells among mNeon cells.

**Extended Data Fig. 13.**
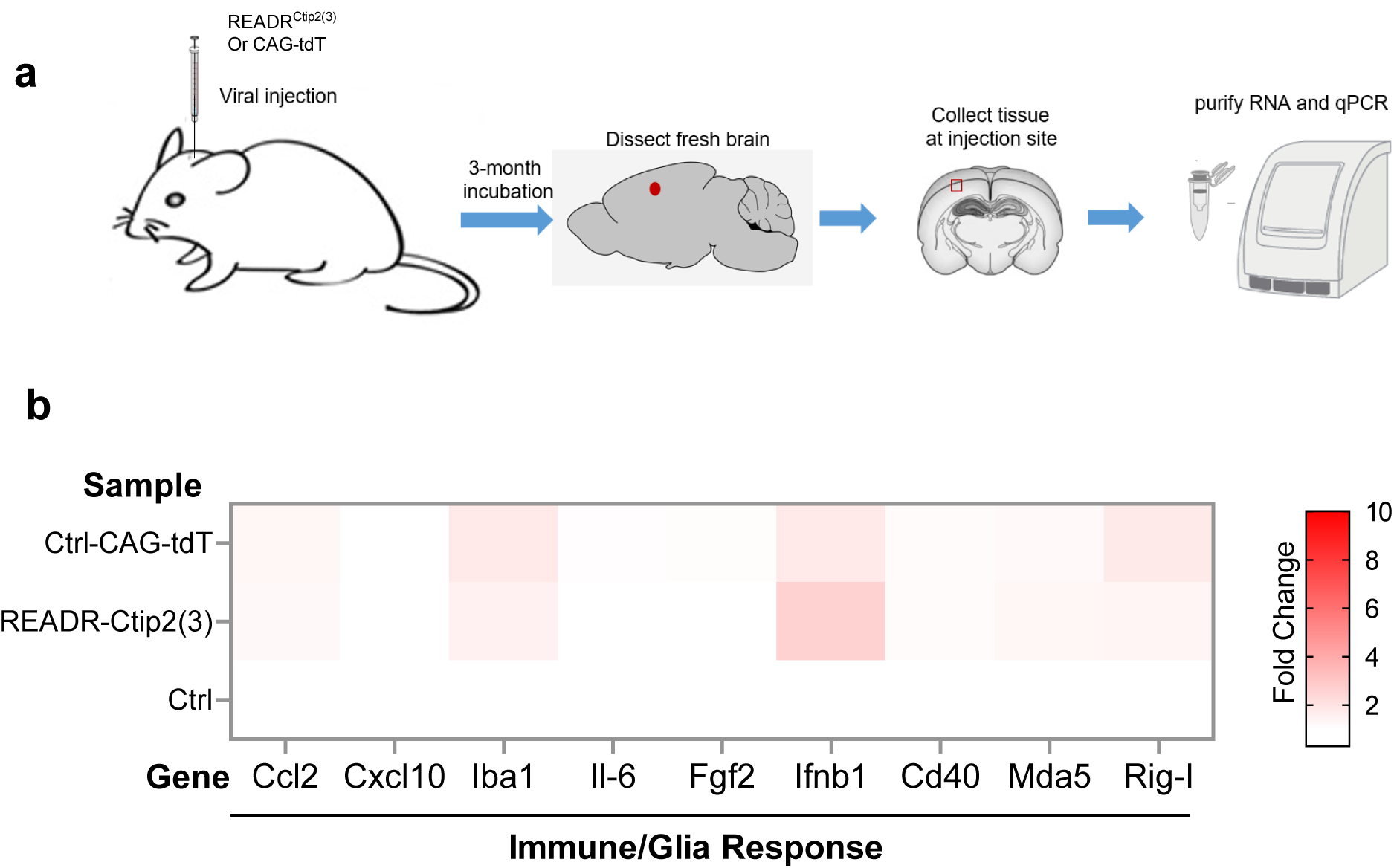
Assessment of cortical cellular immune responses following long-term expression of CellREADR vectors. **a,** Schematic of evaluation of the long-term effects of CellREADR in vivo. For each mouse, *READR^Ctip2(3)^* or *CAG-tdT* control AAVs were injected into S1 cortex and incubate for three months. Fresh brains were dissected and small pieces of cortical tissue at the injection site were collected. Quantitative PCR was performed immediately. S1 tissues of mice without viral injection were used as control. **b,** Heatmap of expression level changes of nine genes implicated in glia activation and immunogenicity.

**Extended Data Fig. 14.**
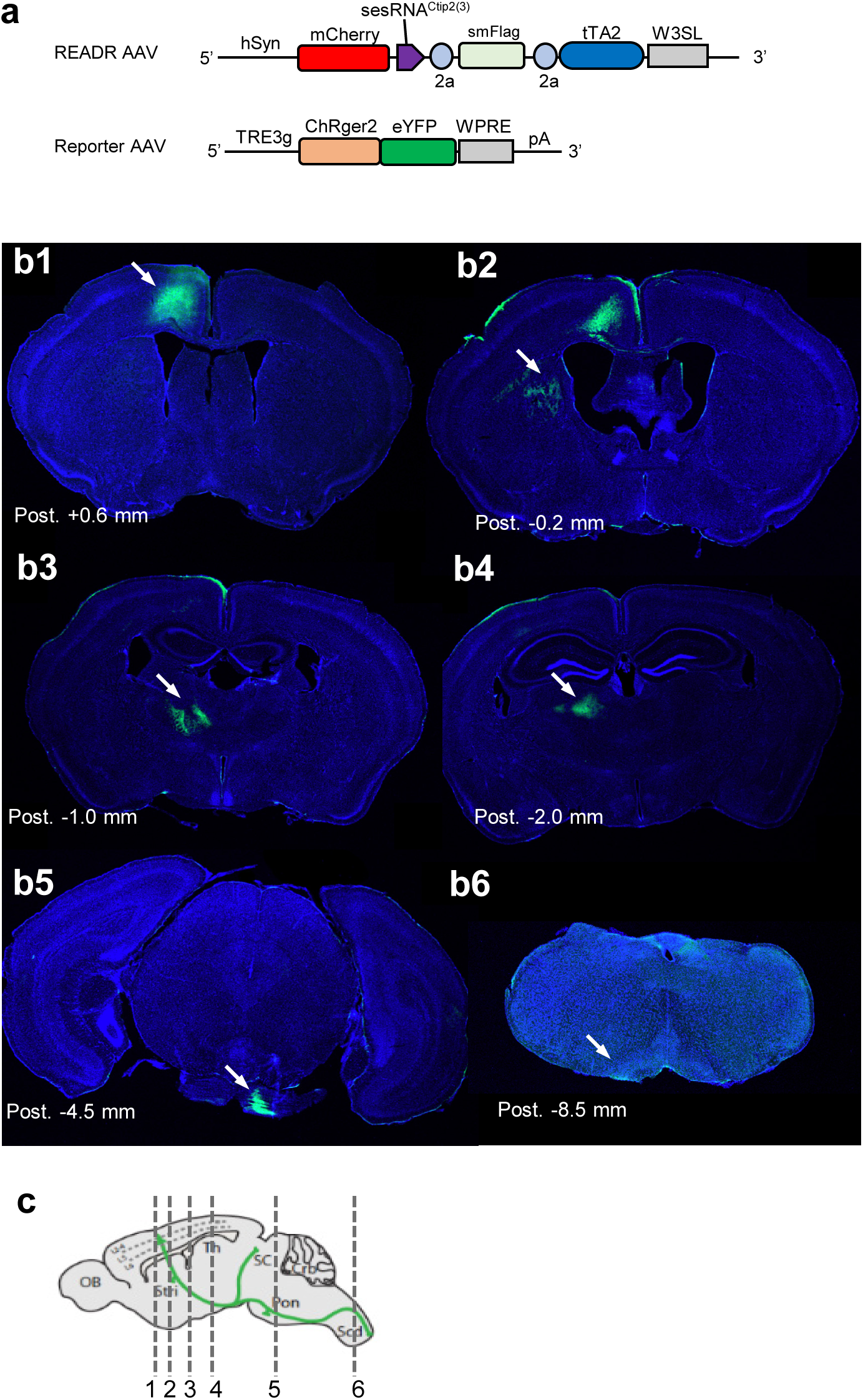
Axonal projection pattern of L5/6 CFPNs in caudal forelimb motor area targeted by AAVs *READR^Ctip2^*and *TRE-ChRger2-eYFP*. **a,** Schematic of binary AAV vectors of *READR^Ctip2(3)-smFlag/tTA2^* and *TRE-ChRger2-eYFP* for optogenetic activation and axonal projection tracing. **b,** Axonal projection pattern of CFPNs infected in CFA. Representative images showing projections to striatum (**b2**), thalamus (**b3, b4**), pons (**b5**) and medulla (**b6**, arrows). **c,** Schematic locations of coronal sections in **b** are shown.

**Extended Data Fig. 15.**
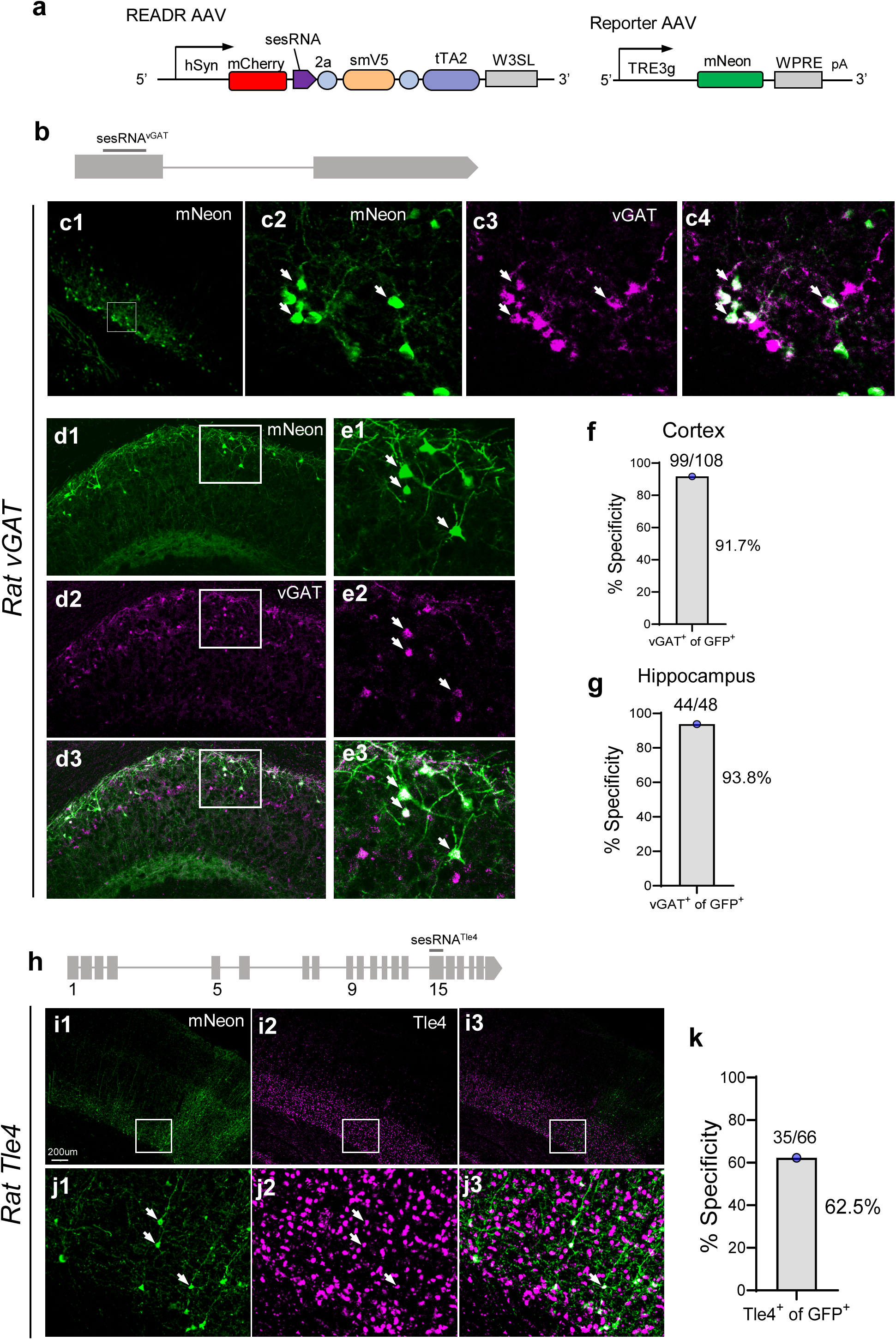
CellREADR targeting of neuron types in rat. **a,** Schematic of binary AAV vectors for cell type targeting in rat. In *READR* vector, a hSyn promoter drives expression of mCherry followed by sequences coding for sesRNA, smFlag and tTA2 effectors. Along with *READR*, a Reporter vector drives mNeonG expression from a TRE promoter in response to tTA2 from the *READR* vector. **b,** Genomic structures of the rat *vGAT* gene with location of a sesRNA as indicated. **c,** AAVs *READR^vGAT^* and *TRE-mNeon* were injected into cortical deep layer and hippocampus. Binary vectors labeled cells shown in cortex (**c1**). **c2,** Magnified view of boxed region in (**c1**). vGAT mRNAs were labeled by in-situ hybridization (**c3**). Co-labeling by mNeon and vGAT mRNA (**c4**). Arrows showed the co-labeled cells. **d,** Cell labeling pattern in the hippocampus CA1 region by co-injection of AAVs *READR^vGAT^* and *TRE-mNeon*. **e,** Magnified view of boxed region in (**d**). Arrows indicate co-labeled cells. **f-g,** Specificity of rat *READR^vGAT^*in rat cortex (**f**) and hippocampus (**g**) measured as the percent of vGAT^+^ cells among mNeon cells. **h,** Genomic structures of the rat *Tle4* gene with the location of a sesRNA as indicated. **i-j,** AAVs *READR^Tle4^* and *TRE-mNeon* (**Fig. 4i**) co-injected into the rat motor cortex labeled cells concentrated in deep layers (**i1**). Tle4^+^ PNs were labeled by TLE4 antibody staining (**i2, 3**). **j,** Magnified view of the boxed region in (**i**). Arrows indicate co-labeled cells by CellREADR and TLE4 antibody. **k,** Specificity of rat *READR^Tle4^*in rat cortex, measured as the percent of TLE4 *^+^* cells among mNeon cells.

**Extended Data Fig. 16.**
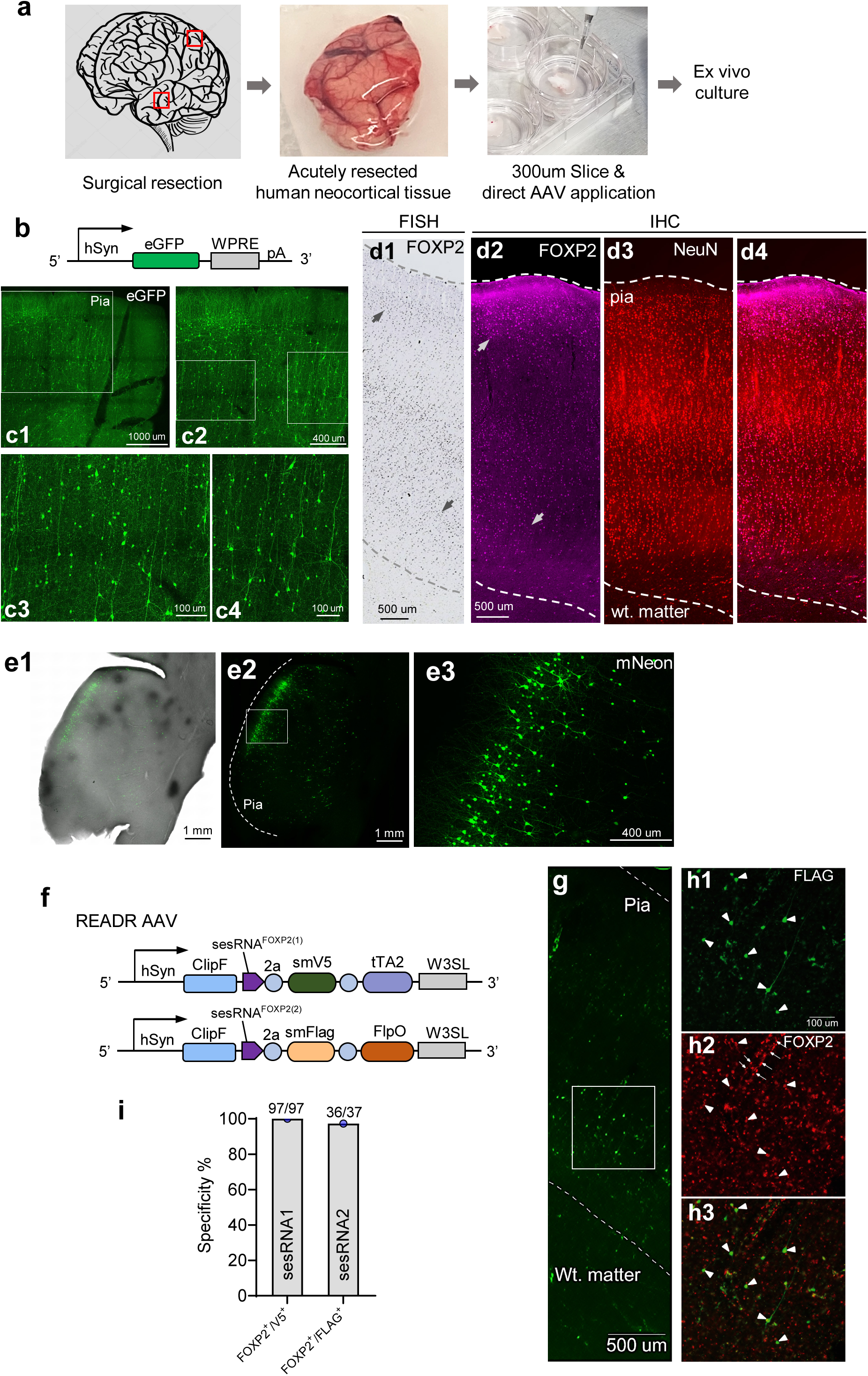
CellREADR vector targeting of neuron types in human cortical *ex vivo* tissues. **a,** Schematic of organotypic platform for of human cortical *ex vivo* tissues. **b,** Schematic of a hSyn-eGFP viral construct used to drive widespread neuronal cell labeling. **c.** *AAVrg-hSyn-eGFP* labeled cells were distributed across all layers, and exhibited diverse morphologies (**c1**). Insets from **c1** (**c2**) and **c2** (**c3, 4**) depict numerous cells with pyramidal morphologies, including prominent vertically oriented apical dendrites. **d,** *FOXP2* expression in human neocortex. *FOXP2* mRNA expression pattern taken from the Allen Institute human brain-map (specimen# 4312), showing upper and deep layer expression (arrows) (**d1**). FOXP2 immunostaining in the current study (magenta) also demonstrated both upper and deep labeling (**d2, 4**). NeuN immunostaining (red) depicting cortical neurons (**d3, 4**). Dashed lines delineate pia and white matter. **e,** *READR^FOXP2(1)^* labeling in an organotypic slice derived from the same tissue used in **d**). Overview of bright field and mNeon native fluorescence in the organotypic slice demonstrating highly restricted labeling, as compared to that observed in **d**. Inset from **e2** (**e3**) depicting morphologies of upper layer pyramidal neurons. **f,** Schematics of two singular vectors of *READR^FOXP2^*. In *READR^FOXP2(1)^*, the hSyn promoter drives an expression cassette encoding ClipF, sesRNA1, smV5, and tTA2. In *READR^FOXP2(2)^*, the hSyn promoter drives an expression cassette encoding ClipF, sesRNA2, smFlag, and FlpO. **g,** Seven days after application of *READR^FOXP2(2)^*AAV on DIV 1, tissue was fixed and stained with antibodies against FOXP2 and FLAG. FLAG-labeled cells from *READR^FOXP2(2)^* exhibited relatively small somata with short apical dendrites (arrowheads). Non-specific background fluorescence signals (e.g. a blood vessel-like profile) are indicated by thin arrows in center panel. **i,** Quantification of CellREADR specificity measured as the percentage of V5^+^ cells (for *READR^FOXP2(1)^*) and FLAG^+^ cells (for *READR^FOXP2(2)^*) labeled by FOXP2 immunostaining, respectively.

**Supplementary Movie 1. Optogenetic activation of mouse injected with binary CellREADR viral vectors**

Optogenetic activation (CFA, 0.5 s) in a mouse injected with binary *READR^Ctip2/3^/Reporter^ChRger2-eYFP^* AAVs induces a stepping forelimb movement. The upward movement involves sequentially, elbow, wrist, and digit flexion followed by extension.

**Supplementary Movie 2. Optogenetic activation of mouse injected with binary PT enhancer viral vectors**

Optogenetic activation (CFA, 0.5 s) in a mouse injected with *PTenhancer/Reporter^ChRger2-eYFP^* AAVs induces elbow extension accompanied by digit extension.

**Extended Data Table. 1.**
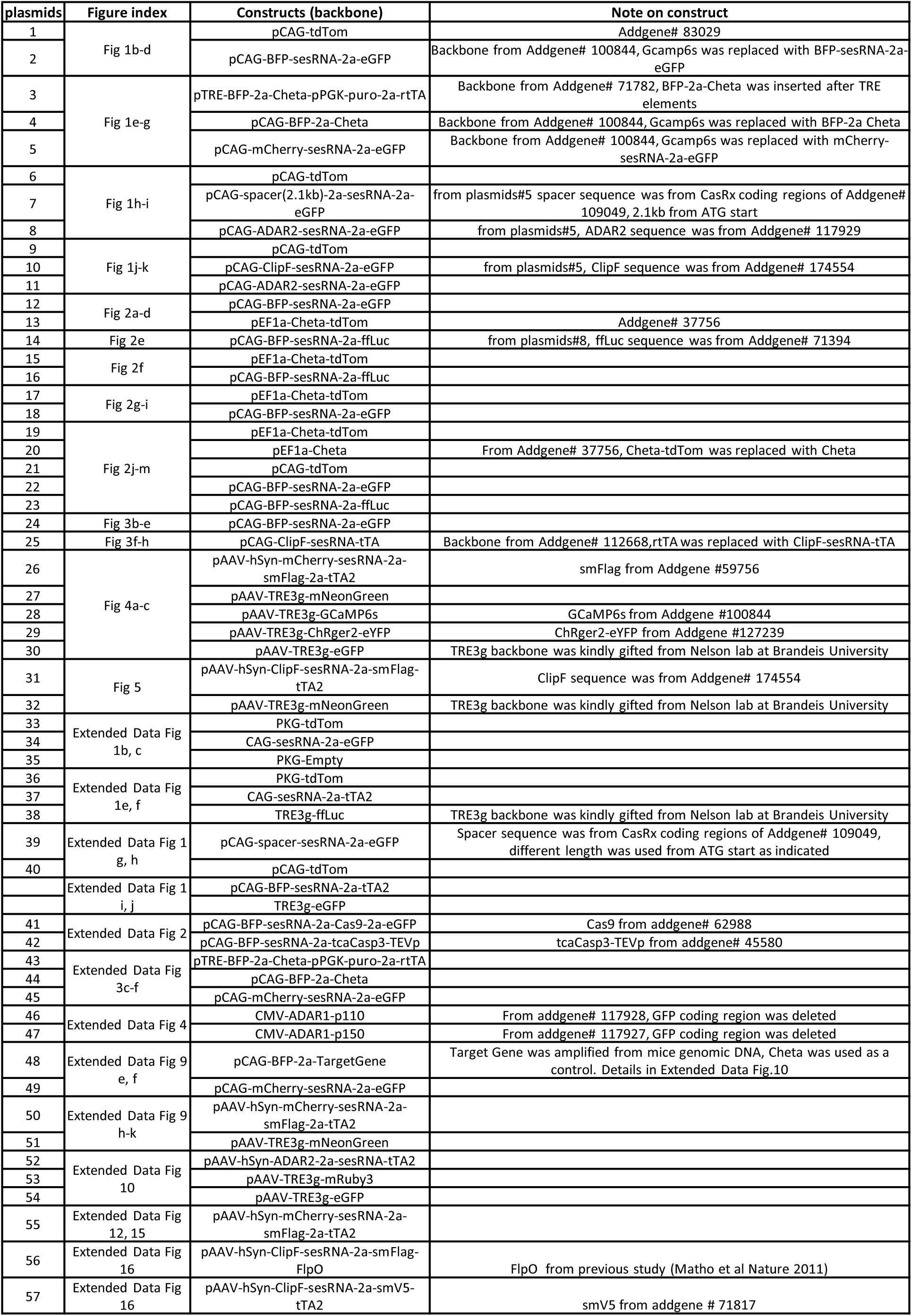
Plasmids used in this study.

**Extended Data Table. 2.**
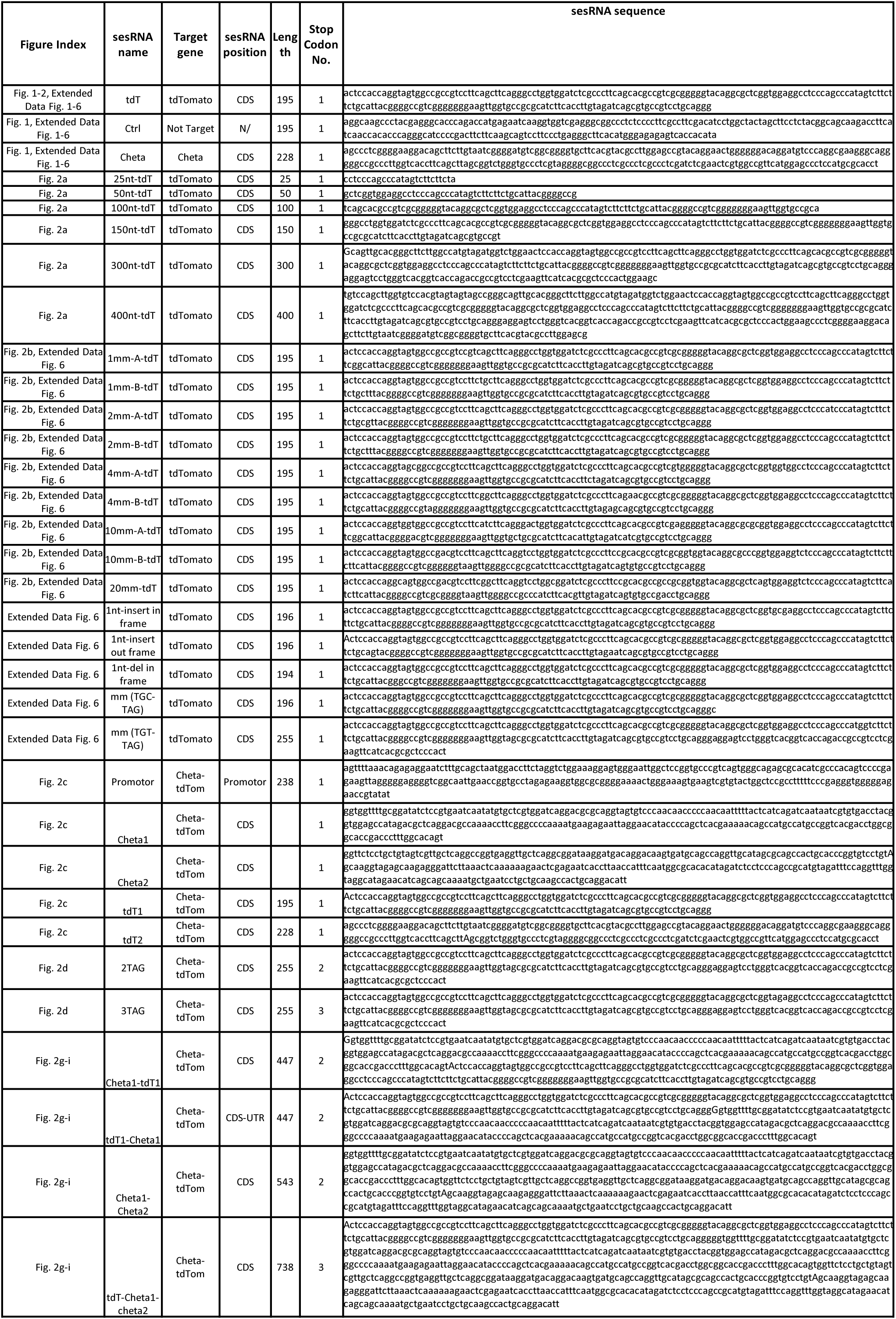
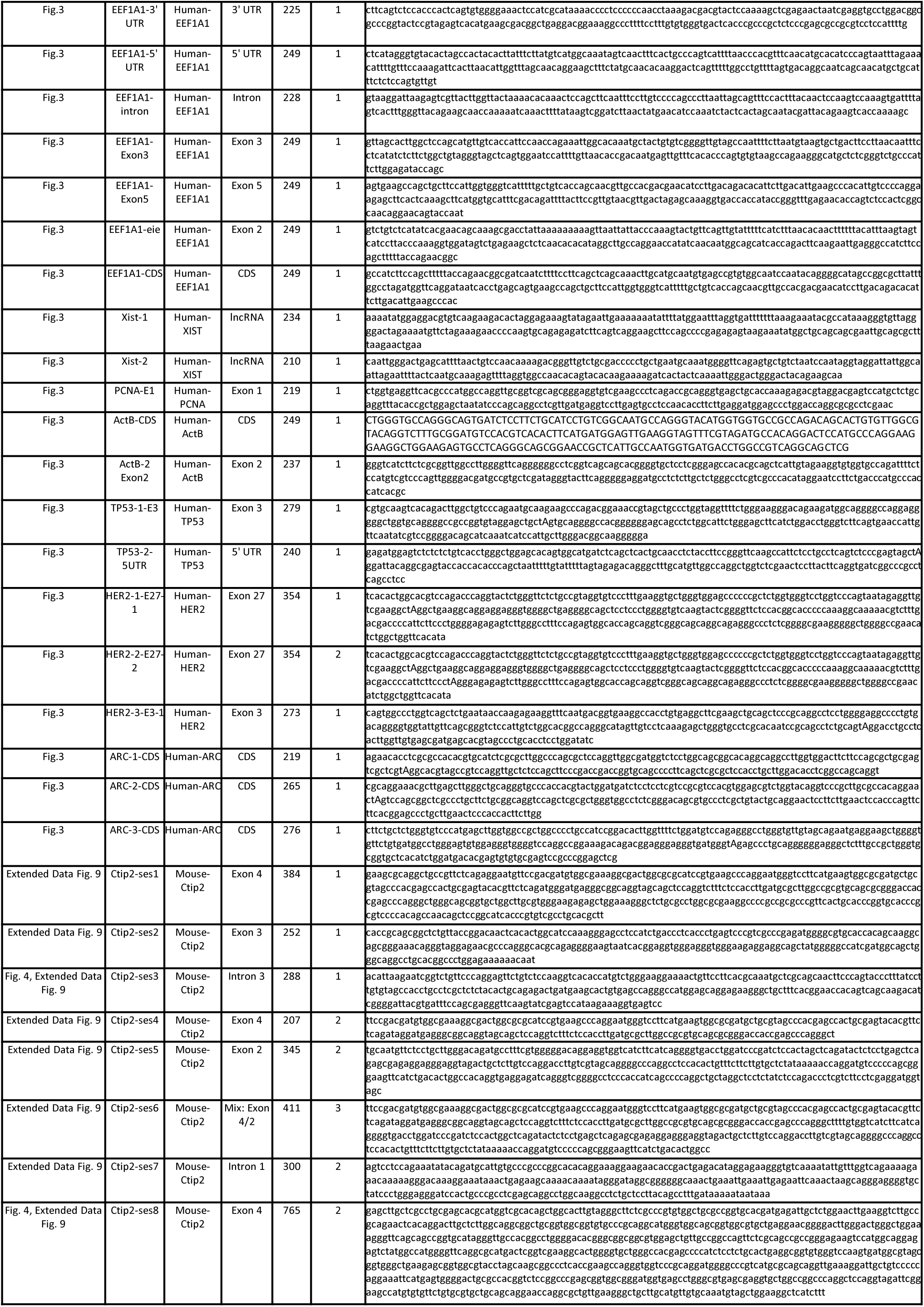
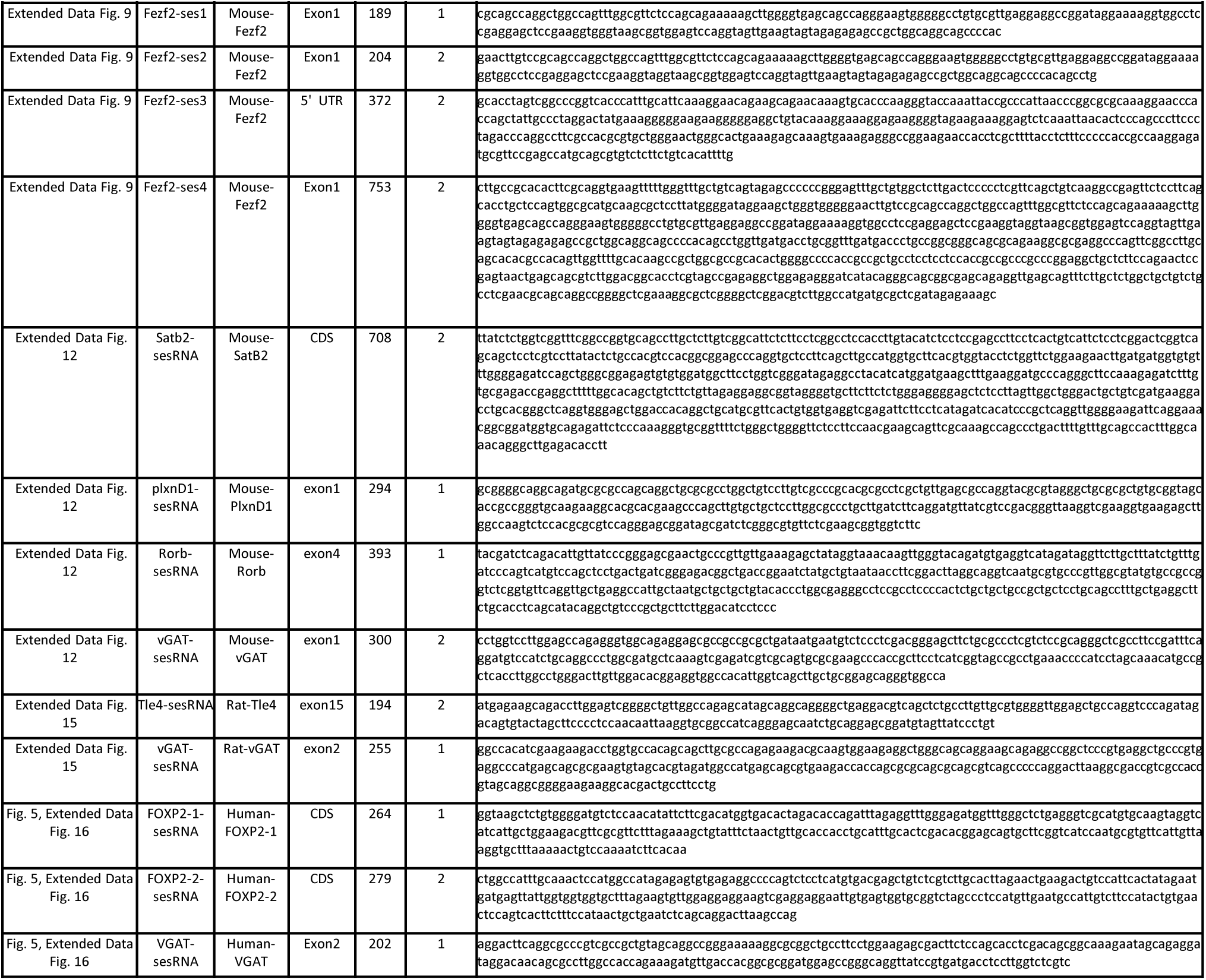
SesRNA used in this study.

**Extended Data Table. 3.**

| **Gene#** | **Gene Name** | **Immune/Interferon Response Related?** | **Fold Change (Log2)** |
| --- | --- | --- | --- |
| OAS2 | 2'-5'-Oligoadenylate Synthetase 2 | Yes | 2.36 |
| OAS1 | 2'-5'-Oligoadenylate Synthetase 1 | Yes | 1.93 |
| APOL6 | Apolipoprotein L6 | No | 1.83 |
| IFI27 | Interferon Alpha Inducible Protein 27 | Yes | 1.82 |
| OAS3 | 2'-5'-Oligoadenylate Synthetase 3 | Yes | 1.81 |
| MX2 | MX Dynamin Like GTPase 2 | Yes | 1.8 |
| XAF1 | XIAP Associated Factor 1 | Yes | 1.75 |
| IFIT3 | Interferon Induced Protein With Tetratricopeptide Repeats 3 | Yes | 1.53 |
| DDX60L | DExD/H-Box 60 Like | Yes | 1.5 |
| IFI44 | Interferon Induced Protein 44 | Yes | 1.49 |
| RSAD2 | Radical S-Adenosyl Methionine Domain Containing 2 | Yes | 1.43 |
| PARP10 | Poly(ADP-Ribose) Polymerase Family Member 10 | No | 1.4 |
| IFIT1 | Interferon Induced Protein With Tetratricopeptide Repeats 1 | Yes | 1.4 |
| TRIM22 | Tripartite Motif Containing 22 | Yes | 1.33 |
| IFI44L | Interferon Induced Protein 44 Like | Yes | 1.29 |
| LRRC37A | Leucine Rich Repeat Containing 37A | No | 1.27 |
| IFIH1 | Interferon Induced With Helicase C Domain 1 | Yes | 1.21 |
| BISPR | BST2 Interferon Stimulated Positive Regulator | Yes | 1.21 |
| IFIT2 | Interferon Induced Protein With Tetratricopeptide Repeats 2 | Yes | 1.21 |
| UBA7 | Ubiquitin Like Modifier Activating Enzyme 7 | No | 1.19 |
| CMPK2 | Cytidine/Uridine Monophosphate Kinase 2 | No | 1.19 |
| ISG15 | ISG15 Ubiquitin Like Modifier | Yes | 1.19 |
| DDX58 | DExD/H-Box Helicase 58 | Yes | 1.19 |
| SAMD9 | Sterile Alpha Motif Domain Containing 9 | No | 1.18 |
| OASL | 2'-5'-Oligoadenylate Synthetase Like | Yes | 1.15 |
| LOC100996724 | PDE4DIP Pseudogene 2 | No | 1.15 |
| DDX60 | DExD/H-Box Helicase 60 | Yes | 1.14 |
| PARP9 | Poly(ADP-Ribose) Polymerase Family Member 9 | Yes | 1.12 |
| MX1 | MX Dynamin Like GTPase 1 | Yes | 1.06 |
| PTBP2 | Polypyrimidine Tract Binding Protein 2 | No | 1.04 |
| IFI6 | Interferon Alpha Inducible Protein 6 | Yes | 1.04 |
| EPSTI1 | Epithelial Stromal Interaction 1 | No | 1.03 |

